# Allergic Asthma Responses Are Dependent on Macrophage Ontogeny

**DOI:** 10.1101/2023.02.16.528861

**Authors:** Robert M. Tighe, Anastasiya Birukova, Yuryi Malakhau, Yoshihiko Kobayashi, Aaron T. Vose, Vidya Chandramohan, Jaime M. Cyphert-Daly, R. Ian Cumming, Helene Fradin Kirshner, Purushothama R. Tata, Jennifer L. Ingram, Michael D. Gunn, Loretta G. Que, Yen-Rei A. Yu

**Affiliations:** Duke University School of Medicine, Department of Pulmonary and Critical Care Medicine, Duke University, Durham, NC; Department of Cell Biology, Duke University School of Medicine, Duke University, Durham, NC; Institute for Life and Medical Sciences, Kyoto University, Kyoto, Japan; Duke University School of Medicine, Department of Neurosurgery, Duke University, Durham, NC

**Keywords:** Macrophages, Asthma, LPS, Ontogeny, Single-Cell RNA-seq

## Abstract

The ontogenetic composition of tissue-resident macrophages following injury, environmental exposure, or experimental depletion can be altered upon re-establishment of homeostasis. However, the impact of altered resident macrophage ontogenetic milieu on subsequent immune responses is poorly understood. Hence, we assessed the effect of macrophage ontogeny alteration following return to homeostasis on subsequent allergic airway responses to house dust mites (HDM). Using lineage tracing, we confirmed alveolar and interstitial macrophage ontogeny and their replacement by bone marrow-derived macrophages following LPS exposure. This alteration in macrophage ontogenetic milieu reduced allergic airway responses to HDM challenge. In addition, we defined a distinct population of resident-derived interstitial macrophages expressing allergic airway disease genes, located adjacent to terminal bronchi, and reduced by prior LPS exposure. These findings support that the ontogenetic milieu of pulmonary macrophages is a central factor in allergic airway responses and has implications for how prior environmental exposures impact subsequent immune responses and the development of allergy.

## Introduction

Macrophages are central mediators of immune responses due to their diverse functional repertoire in homeostasis, initiation, propagation, and resolution of inflammation. In the lung, dysregulation of macrophage functions drives the initiation and progression of several chronic lung diseases, highlighting their role as disease modifiers (1–3). The ability to modulate disease has primarily focused on macrophage responses to polarizing cytokines and/or factors. However, it is becoming clear that macrophage responses depend on their location in tissue structures and cellular origin (4–7). In the lung, tissue-resident macrophages are present in unique tissue compartments (alveolar or interstitial) and have distinct ontogenies (embryonic or bone marrow-derived). In addition, following lung injury, monocytes can be recruited to the lung and differentiate into macrophages (i.e., monocyte-derived macrophages). Research supports that tissue-resident and monocyte-derived macrophages respond differently to injuries (8, 9). Moreover, following the re-establishment of homeostasis, monocyte-derived macrophages can take up niches previously occupied by tissue-resident macrophages and acquire properties resembling tissue-resident macrophages (10, 11). However, there is a limited understanding of how alteration of the ontogenetic composition of tissue-resident pulmonary macrophage by prior injury impacts subsequent immune responses. Given that injury and repair are common features of existence and can change the composition of tissue-resident macrophages (8), understanding how this alters immune responses that lead to lung disease is critical.

The ontogeny of lung macrophages during homeostasis in uninjured rodents has been previously described (12–16). Detailed lineage tracing and parabiosis studies demonstrate that tissue-resident alveolar macrophages are primarily derived from fetal liver monocytes around E12.5. They self-renew without replacement from circulating monocytes and are maintained for the life of the animals (12, 13, 17–19). However, the ontogeny of interstitial macrophages is more complex and less well-defined (20). Resident pulmonary interstitial macrophages are ontologically heterogeneous, consisting of macrophages derived from embryonic and bone marrow-derived progenitors (14). Furthermore, embryonic-derived macrophages appear to differentiate from embryonic precursors (yolk sac vs. fetal liver monocytes) (16). It is presently unclear if yolk sac-derived macrophages are replaced by fetal-liver monocyte-derived cells during development or if embryonic-derived tissue-resident macrophages persist as animals age. Therefore, the clear origin, dynamics, and functions of embryonic-derived interstitial macrophages remain undefined.

Determining pulmonary macrophage complexity has focused on defining their functions in lung injury and repair responses. Following injury and repair, these studies have revealed that the functions of lung macrophages depend, in part, on their origin (i.e. tissue-resident *vs.* recruited monocyte-derived macrophages) (4). In a murine model of bleomycin-induced lung injury and fibrosis, when compared to tissue-resident alveolar macrophages, monocyte-derived alveolar macrophages expressed the majority of genes associated with fibrosis. Deletion of these monocyte-derived macrophages ameliorated fibrosis, supporting a role for monocyte-derived alveolar macrophages as drivers of lung fibrosis (8). Additional studies in lipopolysaccharide (LPS), influenza, and asbestosis-induced lung injury and allergic airway disease models further confirm distinct functional roles for resident vs. monocyte-derived macrophages in the alveolar compartment (9, 11, 21–26). Though less well-defined, different interstitial macrophage subsets also appear to have unique functions during lung injury and repair. In bleomycin-induced lung injury, the loss of Lyve1^hi^MHCII^lo^ interstitial macrophages around blood vessels exacerbated pulmonary fibrosis (5). Additionally, exposure to bacterial CpG DNA increased interstitial macrophages, inhibiting allergic airway responses via interleukin-10 (IL-10) (6) and interstitial macrophages are required for IL-9 mediated allergic airway disease (26). Though these studies clearly support that distinct subsets of alveolar and interstitial macrophages drive different functions in experimental lung diseases, the understanding of how ontogenetic composition of alveolar and interstitial macrophages are altered by prior exposure and the impact of this alteration on subsequent lung immune responses is largely unknown.

The present study defines the ontogeny of embryonic-derived alveolar and interstitial macrophages. We demonstrate that a subset of resident interstitial macrophages are derived from primitive yolk sac macrophages present around E7.5, before the presence of fetal monocytes, which serve as precursors of tissue-resident alveolar macrophages. When undisturbed, these embryonic-derived interstitial macrophages, similar to alveolar macrophages, persist stably through adulthood. We further demonstrate that exposure to LPS causes the replacement of embryonic-derived alveolar and interstitial macrophages with bone marrow-derived macrophages upon re-establishing homeostasis. To explore the consequence of altering resident pulmonary macrophage ontogenetic composition on lung immunity, we exposed these mice to house dust mite (HDM) and assessed allergic airway responses. We identified that prior LPS exposure reduced allergic airway disease, including a reduction in airway hyperresponsiveness, mucus, Th2 cytokine production, and bronchoalveolar lavage and tissue eosinophilia. Using single cell RNA sequencing and lung tissue immunohistochemistry, we identified that allergic gene expression is primarily restricted to a subset of interstitial macrophages present in peribronchiolar lung regions and are tissue-resident derived. This subset was reduced in the LPS pre-exposed mice, supporting the concept that altering pulmonary ontogenetic composition, especially interstitial macrophages, protects from allergic airway disease.

## Materials and Methods

### Experimental animals

C57BL/6, B6.129P2(C)-*Cx3cr1^tm2.1(cre/ERT2)Jung^*/J (Cx3cr1^CreER^) and B6.Cg- *Gt(ROSA)26Sor^tm14(CAG-tdTomato)Hze^*/J (tdTomato) were purchased from the Jackson Laboratory. Male Cx3cr1^CreER^ were crossed with tdTomato mice to generate a Cx3cr1^CreER^/tdTomato lineage reporter. To induce the lineage labeling during the embryonic stage, pregnant females were treated by oral gavage with either 2 mg/mL tamoxifen (Sigma-Aldrich) plus 0.7mg/mL progesterone (Sigma-Aldrich); 3 mg/mL tamoxifen plus 1.4 mg/mL progesterone; or 4 mg/mL tamoxifen plus 2.1 mg/mL progesterone at specified embryonic stage (E7.5 *vs.* E13.5). Due to the impact of tamoxifen on the pregnancy, progesterone was added to aid development and prevent spontaneous abortion. In addition, caesarian section and fostering were performed to allow growth to adulthood. As previously described, bone marrow chimeras were generated using busulfan (27). Experiments were conducted per National Institute of Health guidelines and protocols approved by the Animal Care and Use Committee at Duke University.

### LPS Exposure, Macrophage Depletion, and Allergic Airway Disease Models

LPS (12.5 µg in 50 µL of PBS, Sigma-Aldrich) or PBS was administered by intranasal instillation (i.n.) under light isoflurane anesthesia. This occurred every other week for 3 weeks. Following LPS or PBS exposure, the mice were allowed to recover for >8 weeks before use in the allergic airway disease model using an acute HDM exposure. For the acute model, mice were challenged with 50µl of HDM (100µg total protein, Greer Laboratories Lot#296771, Lenoir, NC) or PBS intranasally (i.n.) on day 1, 7, and 14 under light isoflurane anesthesia. Airway physiology measurements, bronchoalveolar lavage, and tissue harvests were conducted 48h after the last challenge.

### Airway Physiology Measurements

Airway responsiveness to acetyl-β-methacholine (MCh; Sigma-Aldrich) was measured 48h following the final HDM exposure using a computer-controlled small animal ventilator (FlexiVent FX, Scireq, Montreal, Canada). Mice were anesthetized with urethane (1-2 g/kg; Sigma-Aldrich), tracheotomized and intubated with a metal endotracheal tube (18G blunt needle), placed on a 37°C water-heated pad, and mechanically ventilated at a rate of 150 breaths/min with a tidal volume of 10 ml/kg and a positive end-expiratory pressure (PEEP) of 3 cm H_2_O. To block spontaneous respiratory effort, mice were given pancuronium bromide i.p. (0.8 mg/kg; Sigma-Aldrich) five minutes before assessing airway responses. Both 1 second single-frequency and 3 second multi-frequency forced oscillation waveforms were applied using the Flexiware 7.6 software default mouse inhaled dose-response script to measure respiratory mechanics. The resulting pressure, volume, and flow signals were used to determine total respiratory system resistance (R_rs_) and elastance (E_rs_) or Newtonian resistance (Rn), tissue damping (G), and tissue elastance (H) as previously described (28, 29). MCh (10-100 mg/ml) was aerosolized using the FlexiVent FX aeroneb nebulizer attachment. The peak response at each dose was averaged and graphed along with the average peak baseline measurement for each group.

### Bronchoalveolar lavage fluid (BALF) and analysis

Lungs were washed three times with PBS+100µM EDTA+40 µM DTPA via gravity inflation and withdrawn with a syringe inserted into a side port of the tracheal tubing. BALF was spun to remove cells. The BALF was concentrated to 0.25mL using a centrifugal filter (Millipore), and cytokines were measured using a custom Procartaplex (Invitrogen). Cell pellets were treated with RBC lysis buffer (BioLegend) and then re-suspended in 1mL HyClone (Thermo Scientific). Total cells were counted with a Scepter Handheld Cell Counter (Millipore). After staining with Hema 3 solution (Protocol, Kalamazoo, MI), differential cell counts were obtained under light microscopy.

### Flow Cytometry

Flow cytometry was performed on whole lung tissue using our prior published protocols (30, 31). In brief, mice were administered 50uL of 1000/mL heparin subcutaneously before euthanasia. Ten minutes after the injection, the mice were euthanized with isoflurane. The chest wall was opened, and the trachea cannulated. Following cardiac perfusion with 10mL of PBS to remove intravascular red cells, the lung was insufflated past total lung capacity with a digestion solution (1.5mg/ml of Collagenase A (Roche) and 0.4mg/ml DNase I (Roche) in HBSS plus 5% fetal bovine serum and 10mM HEPES). The lung tissue was excised and incubated at 37°C for 30 minutes with intermittent vortexing to generate a cell suspension. After digestion, the samples were washed and strained through a 70uM cell strainer and underwent red cell lysis (ACK RBC lysis solution). The cells were then manually counted and underwent staining for flow cytometry using a previously described flow cytometry panel (Supplemental Table 1 and (30)) of established markers of lung immune cells (32). Flow cytometry was performed on a BD LSRII using BD FACSDiva software and then analyzed using FlowJo version 10 software.

### Histological Analysis

Left lung lobes were isolated and fixed by gravity inflation (20 cm H_2_O) and immersion in 10% formalin. Lungs were paraffin-embedded, cut into 5µm sections, and stained by AML laboratories (Saint Augustine, FL). After individual staining, 10 images of randomly chosen variable size airways were photographed at 10-20x magnification and analyzed in a blinded examination. Lung tissue inflammation was semi-quantitatively determined from hematoxylin and eosin (H&E) and Alcian blue-Periodic acid-Schiff (AB-PAS) stained sections using a four-tiered scoring system as previously described (33, 34). In addition, peri-bronchial fibrosis was determined from Masson’s trichrome staining using Image J (NIH) to calculate the percentage of peribronchial staining within the total tissue area, which included the lumen of the airway.

### Immunofluorescence

Representative unstained formalin-fixed paraffin-embedded (FFPE) sections (5 μm thickness) were stained using automated immunofluorescent techniques on BOND™ RXm Processing Module, utilizing the Bond Research Detection kit (Leica Microsystems) and Opal fluorophores (Akoya Biosciences). In brief, FFPE sections were deparaffinized, hydrated with alcohol, and subjected to heat-induced epitope retrieval with Bond ER1. Slides were washed with Bond wash solution, and non-specific protein binding was blocked with protein block (5 min). For CD206 and Cathepsin K (CTSK) co-staining, the sections were then sequentially incubated with control antibody (Ab) (30 min), Rabbit Polymer (10 min), Opal 520 (10 min), and control Ab (30 min), Rabbit Polymer (10 min), Opal 570 (10 min); or CD206 (E6T5)(Cell Signaling Technology) (0.04 μg/ml) (30 min), Rabbit Polymer (10 min), Opal 520 (10 min), and CTSK (Rabbit anti-mouse CTSK polyclonal antibody)(Novus Biological) (30 min), Rabbit Polymer (10 min), Opal 570 (10 min) Ab fluorophore combination. The nuclei were stained with spectral DAPI (4,6-diamidino-2-phenylindole) solution (Akoya Biosciences). Finally, the sections were cover-slipped using Vectashield HardSet Antifade mounting media (Vector Laboratories). Images were acquired using Zeiss Upright 780 confocal microscope and processed using FUJI/Image J software.

### Microarray Analyses

Yolk sac macrophages (CD45^+^CD11b^+^F4/80^+^) were sorted from the yolk sac layer of E9.5 embryos. AMØs (CD45^+^CD64^+^SSC^hi^CD11b^-^CD11c^+^), IMØs (CD45^+^CD64^+^SSC^hi^CD11b^+^CD11c^-^), Ly6C^+^ classical monocytes (CD45^+^CD64^lo^SSC^lo^CD11b^+^CD11c^-^Ly6C^+^), or Ly6C-non-classical monocytes (CD45^+^CD64^lo^SSC^lo^CD11b^+^CD11c^lo^Ly6C^-^) were sorted from lungs of mice. 10,000 cells were lysed using SuperAmp^TM^ lysis buffer according to manufacturer instructions. Agilent Whole Mouse Genome Oligo Microarrays 8x60K were used to analyze transcriptomic profiles of sorted cells. Microarray processing, including RNA extraction, cDNA synthesis and labeling, hybridization, imaging, and data analysis with normalization were performed by Miltenyi Biotec. Subsequent hierarchical clustering was performed using Partek bioinformatics software.

### Single Cell RNA-sequencing

#### Mapping of reads to transcripts and cells

Lung cells from adult chimeric (CD45.1 *vs.* CD45.2), E13.5 lineage labeled adult Cx3cr1^CreER^/tdTomato mice were sorted into two fractions: Ly6G^-^B220^-^CD45.2^+^ vs. Ly6G^-^B220^-^CD45.1^+^. The presence of tdTomato in CD45.2 fractions was confirmed. For each treatment condition, sorted cells from three animals (n=3) were pooled for library preparation and sequencing. Libraries of cDNA from sorted cells were prepared by Drop-seq platform as previously described by McCarroll Lab (http://mccarrolllab.com/dropseq) and sequenced on Illumina HiSeq 4000 platform. Sequenced reads were mapped to the GRCm38v73 version of the mouse genome and transcriptome (35), binned, and collapsed onto the cell barcodes corresponding to individual cells using the Drop-seq pipeline v2.3.0 (36).

Raw digital expression matrices were generated separately for each of the eight samples, then merged into a single matrix. The Q*uickPerCellQC* function from the R package *scater* (37) v1.18.0 was used to define thresholds for identifying low-quality cells based on QC metrics (UMI count, number of genes detected, percentage of UMIs mapping to the mitochondrial genome). Cells containing >200 genes or <14% mitochondrial UMI were retained for downstream analyses. This filtered data set includes 32,919 cells expressing a total of 22,523 genes. The number of cells per sample ranged from 2,535 (PBS_PBS_CD45.2) to 6,159 (LPS_HDM_CD45.2), with a median of 4,124 cells. We used the R package *Seurat* v4.0.0 and its standard workflow for the integration of multiple samples to integrate the eight samples into a unique dataset (38) before clustering and visualization. Before integration, gene expression was normalized for each sample by scaling the total number of transcripts, multiplying by 10,000, and then log transforming (log-normalization). We then identified the 2,000 genes that were most variable across each sample, controlling for the relationship between mean expression and variance. Next, anchor genes were identified using the *FindIntegrationAnchors* function; then *the IntegrateData function* was used to integrate the eight samples. Integrated data were scaled before performing principal component analyses (PCAs). To distinguish principal components (PCs) for further analyses, the JackStraw method was used to identify 30 statistically significant PCs. Subsequently, a shared nearest neighbor (SNN) modularity optimization-based clustering algorithm was used to determine cell clusters. This was implemented using the *FindNeighbors* function with 30 PCs and the *FindClusters* function with the Louvain algorithm. A range of resolutions was evaluated using the R package *clustree* v0.4.3 to generate a clustering tree. We identified clusters in resolution 0.4 to be particularly stable for higher resolutions. Resolution 0.4 corresponds to 18 cell clusters. Cluster-defining differentially expressed genes (DEGs) were identified using the Seurat function *FindConservedMarkers* for each cluster on the normalized gene expression before sample integration. R package *SingleR* v1.4.0 (39) was used to annotate cell types based on correlation profiles with bulk RNA-seq from the Immunological Genome Project (ImmGen) database (40).

#### Macrophage Subclustering

To examine macrophages, subclustering of macrophages was performed. First, we extracted subsets corresponding to clusters identified as macrophages (26,679 cells). Previously normalized UMI counts were used to identify 2,000 outlier genes on the mean variability plot for each sample; then, the integration of cells originating from each of the eight samples was performed similarly to the whole data set. Thirty statistically significant PCs were identified. Subclustering was then performed on the integrated subsets (resolution = 0.6). A total of 16 macrophage subclusters were identified. Cluster-defining DEGs were again identified using the Seurat function *FindConservedMarkers*. In the macrophage subsets, the three smallest clusters (399, 192, and 21 cells) were identified as small sets of remaining B cells, neutrophils, and mast cells, respectively. As a result, they were excluded from subsequent analyses. To separate the filtered subsets of macrophages into AMØs and IMØs, cells were annotated using the R package *SingleR* with bulk RNA-seq profiles from lung macrophages from Gibbings, Thomas, Atif, McCubbrey, Desch, Danhorn, Leach, Bratton, Henson, Janssen and Jakubzick (41). Cell-type specific markers confirmed the identity of AMØs and IMØs.

#### Subclustering of AMØs and IMØs

The data subset corresponding to AMØs (13,195 cells) was reintegrated and subclustered as described above (resolution of 0.1). Five AMØ subclusters were identified. Differential expression markers characterizing each subcluster were identified using the Seurat function *FindConservedMarkers*.

For IMØs (12,872 cells), a subcluster represented an artifactual cell cluster based on high mitochondrial expression. We, therefore, examined the mitochondrial expression in the IMØ subset for each sample (Figure S4). Five samples display a bimodal distribution of mitochondrial expression that was not detectable at the whole data set level. Thus, the mitochondrial percent expression threshold was lowered to 7.5% for the IMØ subset to filter additional low-quality cells. Reintegration and subclustering on this filtered IMØ subset (10,263 cells) were then performed (resolution of 0.3). None of the resulting seven subclusters identified were technical artifacts.

#### Gene set enrichment and pathway analysis

To further characterize the DEA results in the context of biological pathways, the ‘Core Analysis’ included in Ingenuity Pathway Analysis software (IPA, Ingenuity System Inc, USA, http://www.ingenuity.com) was run on differentially expressed genes (DEGs) for each analysis (p-value < 0.05 and absolute value of log fold-change > 0.2). For both ‘Canonical Pathways’ and ‘Diseases & Functions’ downstream analyses, the p-value is calculated by IPA using a right-tailed

Fisher’s Exact Test. Activation z-score predicts whether the upstream regulator exists in an activated or inactivated state. It is an independently calculated statistical measure of the correlation between relationship direction and gene expression (Z-score> 2 or < -2 is considered significant). The z-score cannot be calculated for all pathways or gene sets: there might be insufficient evidence in the IPA Knowledge Base for confident activity predictions across datasets or having fewer than four analysis-ready molecules in the dataset associated with the pathway.

#### Statistics

Data for animal exposure studies were analyzed using GraphPad Prism 7 software (San Diego, CA). Complex data sets were analyzed using one-way or two-way ANOVA with assessment for multiple comparisons. Data points over 2 standard deviations from the mean were excluded from the data set. The level of significance for all tests was set at *P* < 0.05.

## Results

### Embryonic lineage labeling defines the unique ontogeny of resident pulmonary alveolar and interstitial macrophages

Tissue-resident lung macrophages are derived from embryonic lineages (i.e., primitive yolk sac macrophages or fetal liver monocytes recruited to the lung during embryonic development) or recruited from bone marrow-derived circulating monocytes (i.e. monocyte-derived macrophages) (19, 42, 43). Under homeostatic conditions, these tissue-resident macrophages are maintained through self-renewal for the lifespan or replaced by circulating monocytes (13, 14, 19). The degree and speed of embryonic-derived cell substitution by bone marrow-derived cells may depend on the organ and cellular location (17, 19, 44–46). To address the ontogeny and maintenance of resident pulmonary macrophages, Cx3cr1^CreER^/tdTomato mice were treated with tamoxifen *in utero* to label primitive yolk sac progenies (E7.5) vs. both yolk sac-and fetal liver monocyte-derived progenies (E13.5). Lineage-labeled mice were allowed to age to adulthood to define the ontogeny of macrophages in adult mice (Figure 1A). With E7.5 labeling, approximately 40% of interstitial macrophages (IMØs, CD45^+^CD64^+^CD11b^+^) were tdTomato^+^ (Figure 1B), but 100% of the alveolar macrophage population (AMØ; CD45^+^CD64^+^CD11b^-^) were tdTomato^-^ (Figure 1B). At E13.5, 60% of IMØs and ∼75-80% of AMØs were tdTomato^+^ (Figure 1B), suggesting that IMØs and AMØs have different ontogeny, and a substantial percentage of IMØs are derived from primitive yolk sac progenitors.

**Figure 1.**
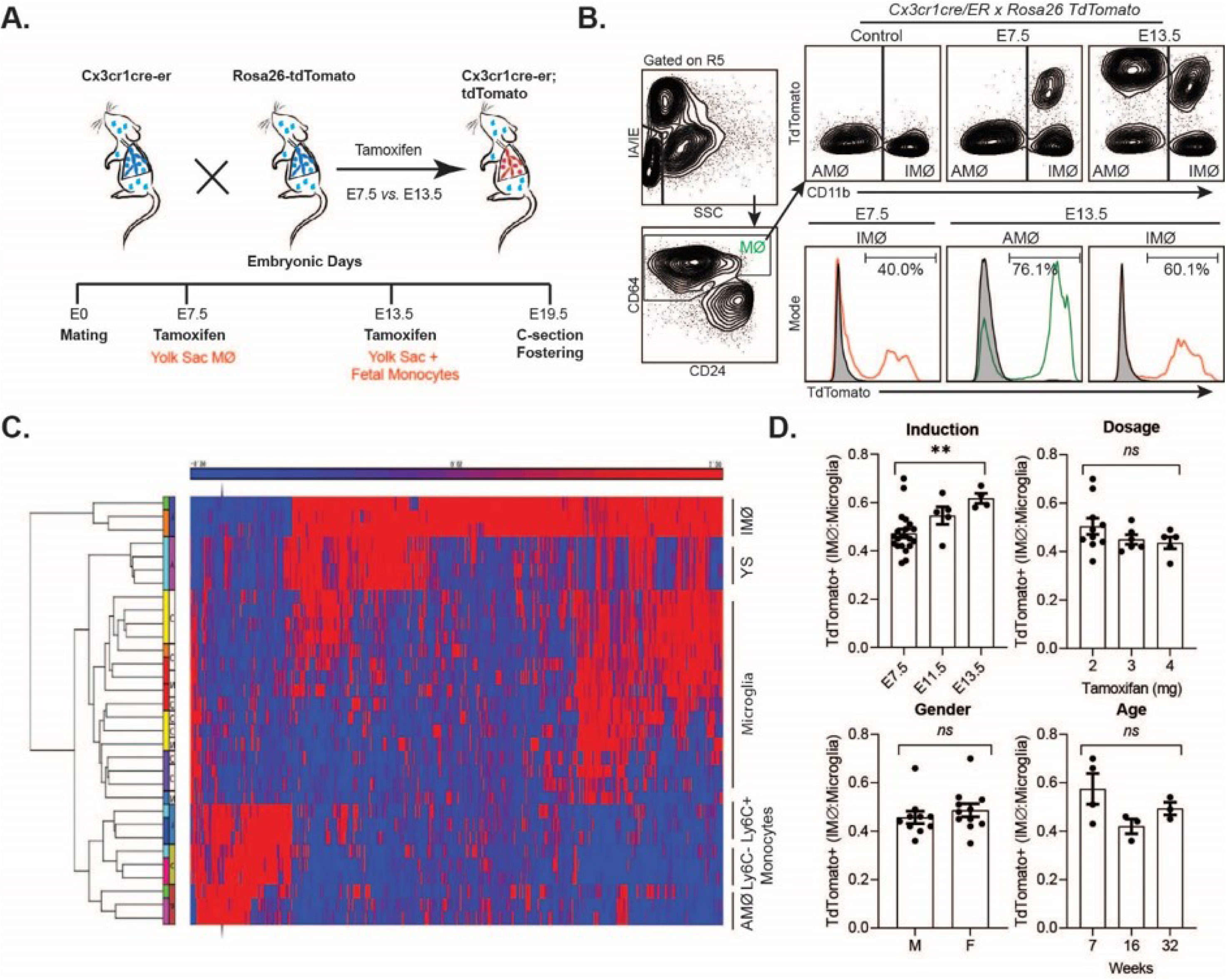
Time-mated lineage labeling of lung macrophages supports that interstitial (IMØ) are Yolk Sac-derived and persist into adulthood. A) Overview of experimental design for time-mated lineage labeling in Cx3cr1cre-er;tdTomato mice induced with tamoxifen at E7.5 to label yolk sac progenitors or at E13.5 to label yolk sac and fetal monocytes. B) Flow cytometry of whole lung tissue in 7 week old mice following embryonic lineage labeling cells in the alveolar macrophage (AMØ) and IMØ compartments based on time of lineage labeling (E7.5 versus E13.5). Live, CD45^+^, Ly6G^-^, CD64^+^ cells were then assessed for TdTomato expression versus CD11b to define AMØ (CD11b^-^) and IMØ (CD11b^+^). Histograms define representative percentages for TdTomato^+^ AMØ or IMØ in adult mice following induction. C) Microarray of sorted AMØ, IMØ, Ly6C^-^ and Ly6C^+^ monocytes, yolk sac (YS) and microglia, reveals clustering of AMØ with monocytes and not IMØ, that have greater overlap with YS. D) Extent of lineage labeling in IMØ, defined as TdTomato^+^ cells in IMØ/microglia, was determined based on the time point of lineage induction, the tamoxifen dose, the sex of mice and the time point of harvest after induction. N=3-10 mice/group, experiments repeated >3 times, **p<0.005, ns=non-significant using ANOVA with post-hoc testing.

In most organs, yolk sac-derived tissue-resident macrophages are diluted or replaced by monocyte-derived macrophages as the animals mature and age (20). To determine the persistence of yolk sac macrophage-derived IMØs and account for labeling efficiency, the percentage of E7.5 lineage-labeled tdTomato^+^ IMØs were normalized to lineage-labeled microglia, as microglia are derived from yolk sac macrophages and persist through the lifespan of animals (47). The ratio of lineage labeled IMØs to microglia remained relatively stable at all ages of animals examined (7, 16, and 32 weeks of age) (Figure 1D-Age). There were no alterations in the ratio based on the dosage of tamoxifen or the gender of the animal (Figure 1D-Dosage and Gender). A minimal increase in tdTomato^+^ IMØ proportion was observed based on tamoxifen-induction timing at E7.5 *vs.* E13.5 (Figure 1D-Induction), which might be due to improved labeling efficiency of IMØs at E7.5 or a minor contribution of the fetal liver monocyte precursors to the IMØ pool. The lineage labeled IMØs to microglia ratio remained relatively stable with different tamoxifen doses, animal gender, and age (Figure 1D-Dosage, Gender, and Age). These findings suggest that, under homeostatic conditions, a substantial portion of tissue-resident pulmonary IMØs, similar to microglia, are derived from yolk-sac progenitors and maintained through local self-renewal. To further confirm the ontogeny of resident pulmonary macrophages, the transcriptome of sorted AMØs, IMØs, microglia, yolk sac macrophages, Ly6C^+^ classical monocytes (Ly6C^+^ Mono), and Ly6C^-^ non-classical monocytes (Ly6C^-^ Mono) were examined by microarray analysis followed by hierarchical clustering (Figure 1C). The AMØs differed from primitive yolk sac progenitors and clustered with monocytes (Ly6C^+^ and Ly6C^-^). The IMØs were distinct from AMØs but shared transcriptomic signatures with primitive yolk sac progenitors. Conflicting reports exist regarding the ontogeny of embryonic-derived IMØs (yolk sac progenitors vs. fetal liver monocytes) (14, 16). However, our findings support that IMØs and AMØs have distinct ontogeny and genetic programming, where AMØs primarily derive from fetal liver monocytes, and a substantial population of IMØs derive from primitive yolk sac progenitors. Additionally, under homeostatic conditions, IMØs derived from embryonic progenitors persist through adulthood.

### Resident macrophage ontogeny regulates allergic airway responses

When the tissue-resident macrophage niche is perturbed through experimental depletion or inflammation, monocyte-derived macrophages acquire a tissue-resident phenotype with self-renewal properties (10, 15, 44, 45, 48, 49). After re-establishing homeostasis following environmental perturbation, monocyte-derived macrophages often closely resemble those of embryonic-derived resident macrophages (10, 49). However, it is unclear whether alteration of tissue-resident macrophage ontogenetic composition impact immune response to subsequent insults. To address the contribution of altered macrophage ontogenetic composition to lung immune responses, we induced the replacement of embryonic-derived by monocyte-derived macrophages. Following re-establishment of homeostasis, we assessed the impact of the embryonic to monocytic-macrophage switch on allergic airway responses to HDM (Figure 2A). Macrophage ontogenetic alteration was induced by *i.n.* exposure to LPS. The mice recovered for > 8 weeks to allow complete resolution of LPS-induced acute inflammation. Resolution of inflammation was assessed by histologic analyses (in Figure 2D and S2) and immunophenotyping by flow cytometry (Figure S3) where the PBS and LPS exposed mice did not show significant differences before HDM exposure. To assess the effectiveness and persistence of replacement at >8 weeks after LPS exposure, we determined the change in the proportion of tdTomato^+^ AMØs and IMØs in E13.5 embryonically-labeled Cx3cr1^CreER^/tdTomato mice. Compared to PBS control mice, LPS-exposed mice had an approximate 50% reduction in lineage labeled tdTomato^+^ AMØs or IMØs, supporting the recruitment and persistence of monocyte-derived macrophages (Figure 2B). After confirmation of macrophage ontogenetic composition alteration, we assessed whether this alteration impacted allergic responses to the HDM challenge. We observed that mice exposed to LPS and then HDM (LPS+HDM, red open box), when compared to mice exposed to PBS and then HDM (PBS+HDM, red closed box), had reduced airway physiologic responsiveness, including resistance and elastance (Figure 2C). While macrophage alteration did not impact the degree of total inflammatory cell infiltrate (H&E) or peri-bronchial fibrosis (Mean % Trichrome), replacement by monocyte-derived macrophages in the setting of repeated LPS exposure led to reduced mucus accumulation in response of HDM exposure (Mean PAS score) (Figure 2D) Even though, alteration in macrophage composition did not impact overall HDM-induced inflammatory cell infiltration (H&E), in LPS+HDM animals, there was a significant reduction in eosinophil numbers in bronchoalveolar fluid (BALF) and lung tissue (Figure 2E). Consistent with decreased eosinophils, there was a reduction in BAL T helper 2 (Th2) cytokines, including IL-4, IL-5, and IL-13, which are known to modulate eosinophil trafficking and functions (Figure 2F). This data supports that prior LPS exposure, which alters the ontogenetic composition of AMØs and IMØs, improves allergic airway responses to HDM.

**Figure 2.**
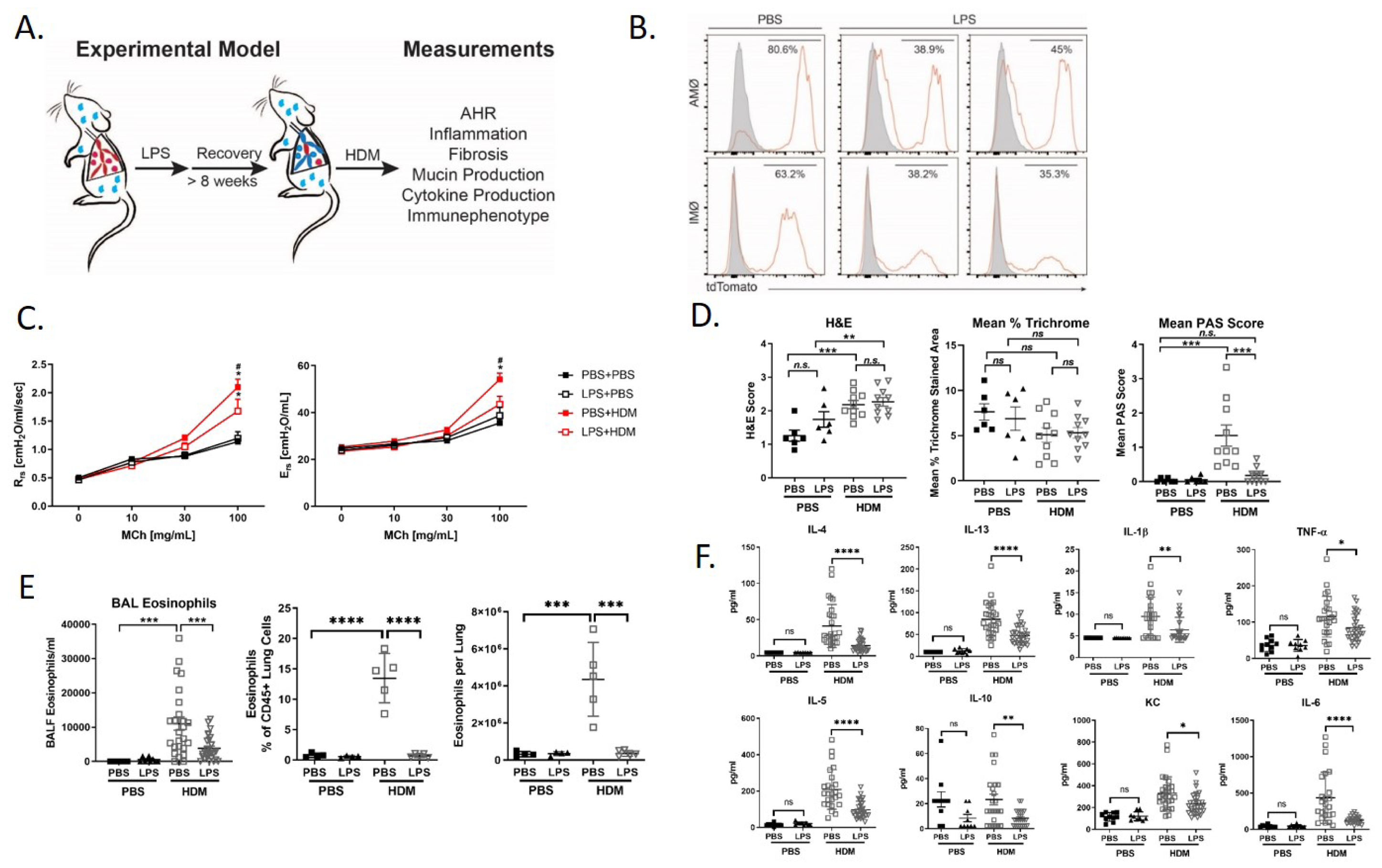
Replacement of embryonic-derived macrophages with bone-marrow derived macrophages with LPS exposure and recover attenuates house dust mite (HDM) allergic airways responses. A) Overview of experimental design to assess the impact of macrophage turnover on allergic airway responses. Mice underwent intranasal (i.n) LPS exposure (12.5µg) every other week for 3 doses. They were then allowed to recover for > 8 weeks. Following recovery, mice underwent acute HDM exposure and were assessed for phenotypic allergic airway responses. B) Cx3cr1^CreER^/tdTomato mice underwent lineage reporter induction with tamoxifen at E13.5 and then allowed to age till 6 weeks. At 6w, i.n. LPS or PBS exposure as described above was performed. Following >8 weeks of recovery from the exposure, lung tissues were harvested and processed for flow cytometry to define tdTomato expression in AMØ and IMØs. LPS causes turnover of embryonic-derived macrophages to bone-marrow derived macrophages as defined by a reduction in the percentage of TdTomato+ cells. C-F) LPS mediated turnover reduces HDM-mediated allergic airway responses including reduced airway hyperresponsiveness to increasing doses of methacholine (C, R_rs_-resistance, E_rs_-elastance), reduced periodic acid–Schiff (PAS) staining (measure of mucus) without a difference in hematoxylin and eosin (H+E) or trichrome staining (D), decreased lung tissue (as a % of CD45+ cells or total cells) and bronchoalveolar lavage fluid (BALF) eosinophils (E), and decreased BALF cytokines including IL-4, IL-13, IL-1β, TNF-α, IL-5, IL-10, KC and IL-6 (F). n=10 mice/group in the PBS/PBS and LPS/PBS groups and n=30 mice/group in the PBS/HDM and LPS/HDM groups. *p<0.05 when compared to PBS control or ^#^p<0.05 when compared between PBS-HDM and LPS-HDM groups by 2-way ANOVA with testing for multiple comparisons. *p<0.05, **p<0.005, ***p<0.0005, ****p<0.00005 for other comparisons by 1-way ANOVA or Students T-test, n.s.=non-significant.

### Single-cell RNA sequencing reveals a unique lung resident IMØ subset that regulates HDM responses

To determine the impact of LPS pre-exposure and macrophage ontologic alterations on HDM allergic responses, we performed single cell RNA-sequencing (ScRNAseq) to define immune cell composition and gene expression. To define pulmonary macrophage ontogeny, we generated bone marrow chimera by transferring CD45.1 bone marrow donor cells into busulfan-treated CD45.2, embryonic labeled (at E13.5) Cx3cr1^CreER^/tdTomato recipient mice (Figure 3A). We have previously shown that busulfan treatment induces efficient bone marrow myeloid ablation while having minimal effects on resident pulmonary macrophages (10, 27). Thus, we can clearly distinguish tissue-resident macrophages (CD45.2^+^tdTomato^+^) from those arising from the bone marrow compartment (CD45.1^+^tdTomato^-^). Following engraftment, these mice were exposed to PBS or LPS, allowed to recover for >8 weeks, and exposed to PBS or HDM. Following exposure, lungs were harvested and digested to generate a single cell suspension. To enrich for myeloid cells and to segregate tissue-resident and hematopoietic cells, the lung single cell suspensions were sorted for live, CD45^+^ Ly6G^-^ B220^-^ cells and then separated by CD45.1 vs. CD45.2 (Figure 3A). ScRNAseq of sorted cells identified distinct populations of cells based on a UMAP plot, and annotated by cell type-associated markers. Clustering identified expected clusters of macrophages (*Trem2* and *Cxcl2*), T cells (*Cd3e* and *Cd3g*), dendritic cells (*Xcr1* and *Fscn1*), and monocytes (*Cx3cr1*) (Figure 3B). There were small clusters of residual endothelial cells, B cells, and epithelial cells (Figure 3B). To define if there are unique clusters based on exposure condition and ontogeny, macrophages were subclustered into AMØs (*Itgax* and *Car4*) and IMØs (*Itgam*, *Csf1r*, *Cx3cr1*, *Ccr2*, and *Lyve1*) groupings (Figure 3C) as well as subclustered based on both exposure conditions (PBS *vs*. LPS, and PBS *vs.* HDM) and ontogeny (CD45.1 *vs.* CD45.2)(Figure 3D). We observed an expansion of IMs in response to HDM exposure and a specific cluster of cells (Figure 3D, dark blue) arising primarily from CD45.2 resident pulmonary interstitial macrophages in the setting of HDM challenge. These observations suggest the unique macrophage clustering based on ontogeny and exposure condition.

**Figure 3.**
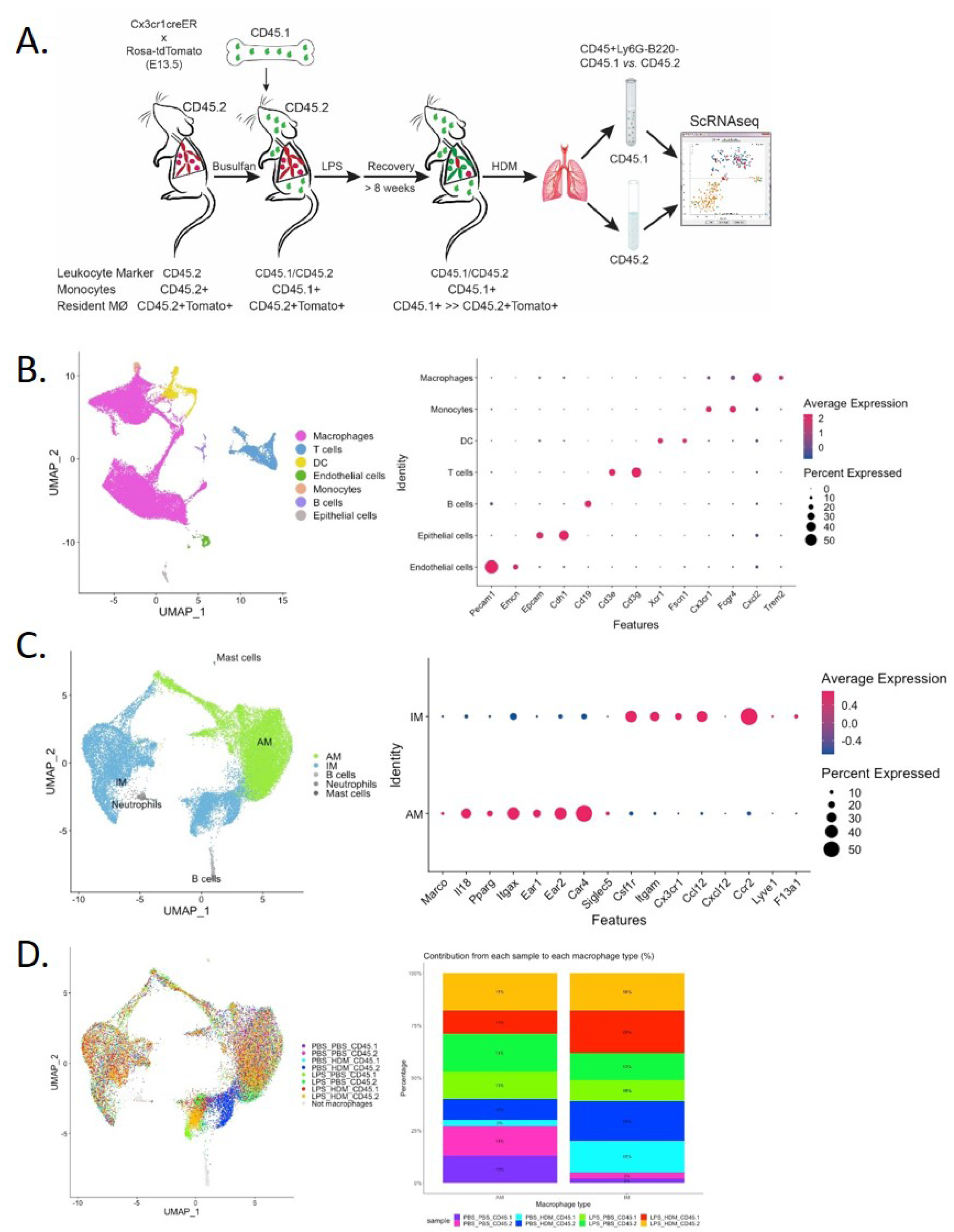
Single cell RNA sequencing of immune cells allows segregation of alveolar and interstitial macrophages and identifies unique clustering based on exposure condition. A) Bone marrow chimeras were generated from lineage labeled Cx3cr1^CreER^/tdTomato mice by injecting CD45.1 donor cells into CD45.2 recipients to define resident (CD45.2) and recruited (CD45.1) cell populations. Mice were exposed in the following groupings PBS_PBS, PBS_HDM, LPS_PBS and LPS_HDM. Following exposures, lungs were harvested and processed for sorting. Live, Ly6G^-^, B220^-^ cells were segregate based on CD45.1 from CD45.2 expression and processed for single cell RNA-seq. B) Clustering of immune cells based on defined markers identifies macrophages (pink), dendritic cells (DC, yellow), monocytes (orange), and T cells (blue). C) Sub-clustering of macrophages segregates interstitial (IM, blue) from alveolar macrophages (AM, green) based on annotation using established markers. D) Display of sub-clustering of macrophages based on exposure condition and whether the cells are recruited (CD45.1) or tissue-resident (CD45.2) identifies distinct clustering based on exposure and origin. Bar graph reveals that IMs increase as a percentage in response to exposures when compared with the PBS_PBS or LPS_PBS groups. Data was pooled from individual sorted mouse lung samples (n=3 mice per exposure group) following the flow sorting.

### Differential effects of ontogeny on alveolar and interstitial macrophages in allergic airway responses

Both AMs and IMs have been implicated in regulating allergic asthma responses; thus, we performed further analyses of both AMØs and IMØs to determine how altering ontogenetic milieu affects their transcriptomic profiles. Sub-cluster analysis of AMs revealed 5 unique clusters (AM1-5; Figure 4A-B). Based on differential gene expression and Ingenuity pathway (IPA) analysis (Figure S4 and S5A), AM1 consisted primarily of cells that down-regulated pro-inflammatory and proliferative processes (*e.g.,* interferon signaling, chemokine signaling, neuroinflammatory signaling, mitosis, metaphase signaling, and cell cycle control of chromosome replication); AM2 represent a cluster of proliferating cells (*e.g.,* Mki67; DNA replication, mitosis, and cell cycle progression); and AM3-5 participated in immune responses (*e.g.,* myeloid cell activation, chemotaxis/migration/accumulation, phagocytosis, or systemic autoimmune syndrome). Overlaying clusters based on exposure conditions and ontogeny did not reveal unique condition/ontogeny-specific clustering either by UMAP or the individual cell/cluster percentages (Figure 4C). No prominent contribution of macrophage ontogeny (CD45.2 vs. CD45.1) was observed to HDM responses (Figure S4); however, differences between the pre-HDM exposure conditions (PBS vs. LPS pre-exposure) were observed.This data largely suggests that LPS pre-exposure, but not ontogeny, modulates AMØs phenotype in response to HDM challenge.

**Figure 4.**
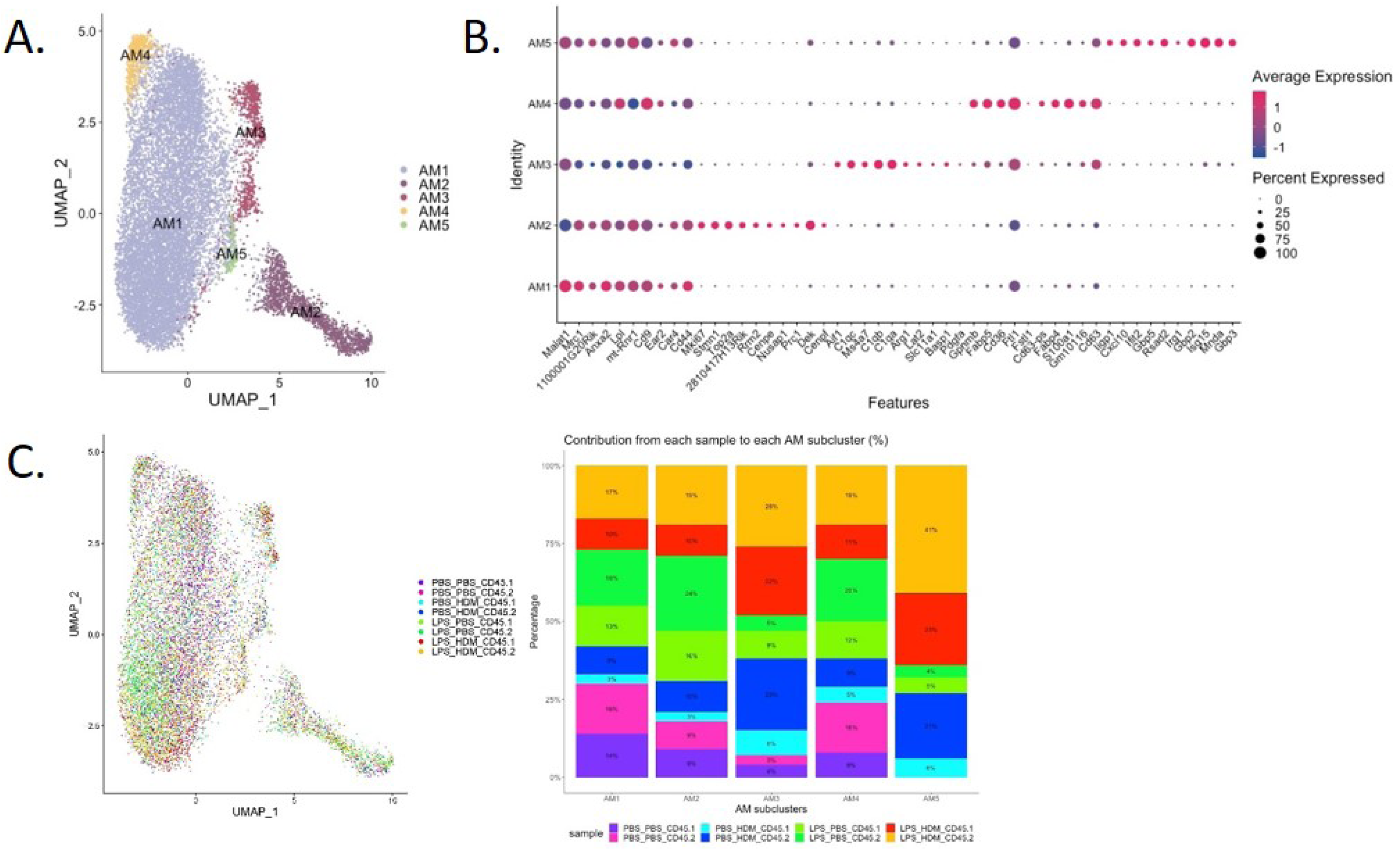
Alveolar macrophage sub-clustering reveals unique clusters with distinct gene expression patterns but without evidence of distinct clustering based on ontogeny. A) Alveolar macrophages (AM) are sub-clustered, identifying 5 unique clusters (AM1-5) defined by individual gene expression patterns (B). C) An overlay of the clusters based on exposure and CD45.1 (bone-marrow derived) or CD45.2 (tissue-resident) group in UMAP plot; as well as bar graph depicting or percentage contribution by each sample to individual AM subclusters. Data do not support clear cluster segregation based on ontogeny. Data was pooled from individual sorted mouse lung samples (n=3 mice per exposure group).

Analyses of IMØs identified 7 distinct clusters (Figure 5A). Similar to AMØ clusters, differential gene expression, and IPA analyses define cells that down-regulated proliferative pathways (IM1), upregulated proliferation pathways (IM5), and cells involved in overlapping inflammatory pathways (IM3, 4, 6, and 7; *e.g.,* iNOS, IL-1, IL-6, and Trem-1 pathways) (Figure 5A + Figure S5B and data not shown). Interestingly, IM2 consisted mainly of the population of previously observed cells (dark blue) that arise primarily from CD45.2 resident pulmonary macrophages following the HDM challenge (CD45.2 PBS_HDM, blue, Figures 3D and 5B-C). This response was largely not observed in animals re-challenged with PBS regardless of ontogeny or prior-LPS exposure (CD45.1 PBS_PBS, CD45.2 PBS_PBS, CD45.1 LPB_PBS, and CD45.2 LPS_PBS) and animals pre-exposed to LPS followed by HDM challenge regardless of macrophage ontogeny (CD45.1 or CD45.2 LPS_HDM) as there was a minimal cellular contribution to the unique IM2 sub-cluster (dark blue cells) by these samples (Figure 5C). Additionally, top gene analysis identified that the IM2 cluster was defined by expression of the *Chi3l3* (chitinase-like 3/Ym1), which has been previously associated with Th2, eosinophilic, and allergic airway responses (Figure 5D) (50–53). Additionally, the *Chi3l3* gene expression level was most prominent in the CD45.2 PBS_HDM within the IM2 subcluster (*Chi3l3*^hi^; dark blue cells) (Data not shown). These findings suggest that *Chi3l3*^hi^ CD45.2 IM2 cells, uniquely and dynamically increased following HDM exposure, have the potential to regulate allergic airway responses. Therefore, as opposed to AMØs, IMØs ontogenetic composition appears to be a regulating factor in allergic airway responses. Furthermore, replacing embryonic-derived resident interstitial pulmonary macrophages with monocyte-macrophages dampens this HDM-induced allergic inflammation.

**Figure 5.**
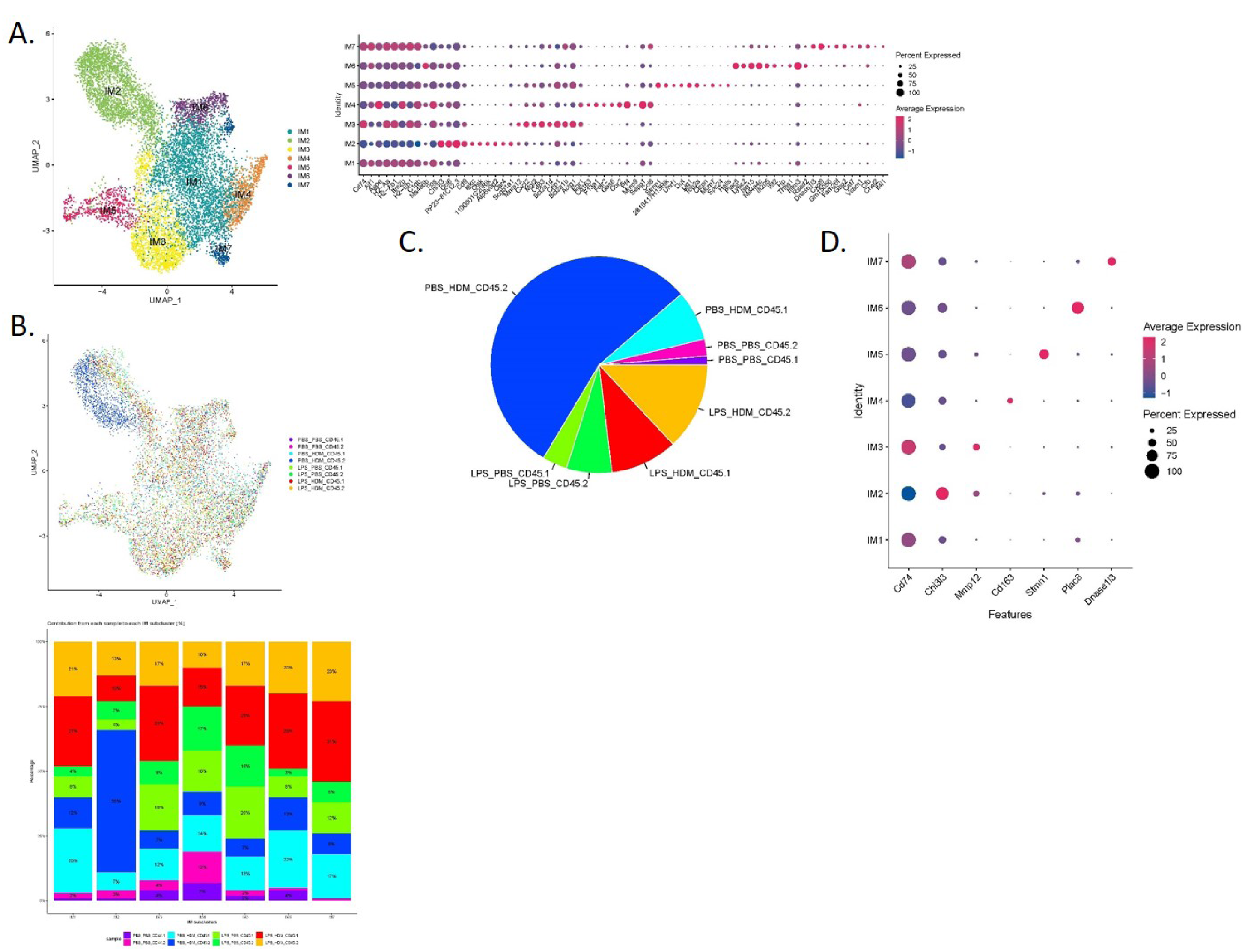
Interstitial macrophage sub-clustering reveals a unique IMØ subcluster that segregates by exposure condition, ontogeny, and is defined by expression of Chi313. A) Sub-clustering of interstitial macrophages (IM) defined by individual cellular markers, identifies seven unique clusters (IM1-7). B) Overlay of IM clusters on UMAP and bar graph based on exposure condition and cell ontogeny identifies that IM2 cluster segregates based on CD45.2 (resident origins) and HDM exposure, C) Pie chart demonstrating the distribution of the IM2 cluster based on the exposure condition and cell origins. This highlights that tissue-resident CD45.2 IMØ from the PBS_HDM exposed mice as the predominant constituent of the IM2 cluster. D) Top gene expression for each individual cluster highlights that cluster IM2 is defined by gene expression for Chi313. Data was pooled from individual sorted mouse lung samples (n=3 mice per exposure group) following the flow sorting.

### Allergic response-associated pathways are enriched in resident-derived interstitial macrophages

To better define the genetic pathways defining resident macrophages in the IM2 cluster, we performed an analysis based on ontogeny and exposure. A heatmap of top differentially expressed genes based on either the PBS_HDM or the LPS_HDM exposures and lineage (CD45.1 vs. CD45.2) of the IM2 cluster was generated (Figure 6A). This dataset revealed clear separation in the gene expression pattern in the CD45.2 PBS_HDM cells from other conditions (CD45.1 PBS_HDM, and CD45.1 *vs.* CD45.2 LPS_HDM). To clarify the pathways involved in this response, we performed an IPA pathway analysis to assess the potential functions of CD45.2 PBS_HDM IM2 cells. We focused on genes and pathways related to macrophage functions that could regulate allergic asthma, including processes related to eosinophil signaling, inflammation, vascular-associated signals, remodeling, and metabolism/death. Consistent with high *Chi3l3* expression, the CD45.2 PBS_HDM cells in IM2 were enriched for many macrophage-mediated inflammatory and remodeling processes relevant to allergic airway responses. Notably, these included those that specifically regulate eosinophilic responses (e.g., CCR3 signaling in eosinophils, Fc Epsilon RI signaling, and IL-3) (Figure 6B). These pathways were down-regulated in the LPS_HDM cell groups. In addition, we observed that CD45.1 monocyte-derive macrophages (CD45.1_PBS_HDM) exhibited reduced allergic pathway activation compared to CD45.2 tissue-resident macrophages (CD45.2_PBS_HDM), suggesting that replacement of CD45.2 tissue-resident macrophages by CD45.1 monocyte-derived macrophages reduces macrophage-derived allergic responses. While to a lesser degree than the difference observed across different ontogeny (CD45.2 *vs.* CD45.1), we also observed a reduction of allergic pathway signaling in CD45.2 tissue-resident macrophages following LPS exposure (CD45.2_LPS_HDM) when compared to PBS exposed CD45.2 tissue-resident macrophages (CD45.2_PBS_HDM) suggesting a decrease of tissue-resident macrophage allergic potential in addition to replacement by CD45.1 cells (Figure 6B). These findings further indicate that there may be effects related to replacement by CD45.1 cells and the impact of prior exposure on the allergic potential of tissue-resident macrophages. We also observe that LPS pre-exposure leads to the down-regulation of small numbers of inflammatory pathways (*i.e.,* CCR5, IL-8, and thrombopoietin) in a lineage non-specific fashion (Figure S6). And, as expected, there were upregulated pathways in response to HDM, which were shared across ontogeny or pre-exposure conditions (Figure S6). When taken together, these findings support that the ontogeny of interstitial macrophages is important for subsequent allergic responses to an allergen.

**Figure 6.**
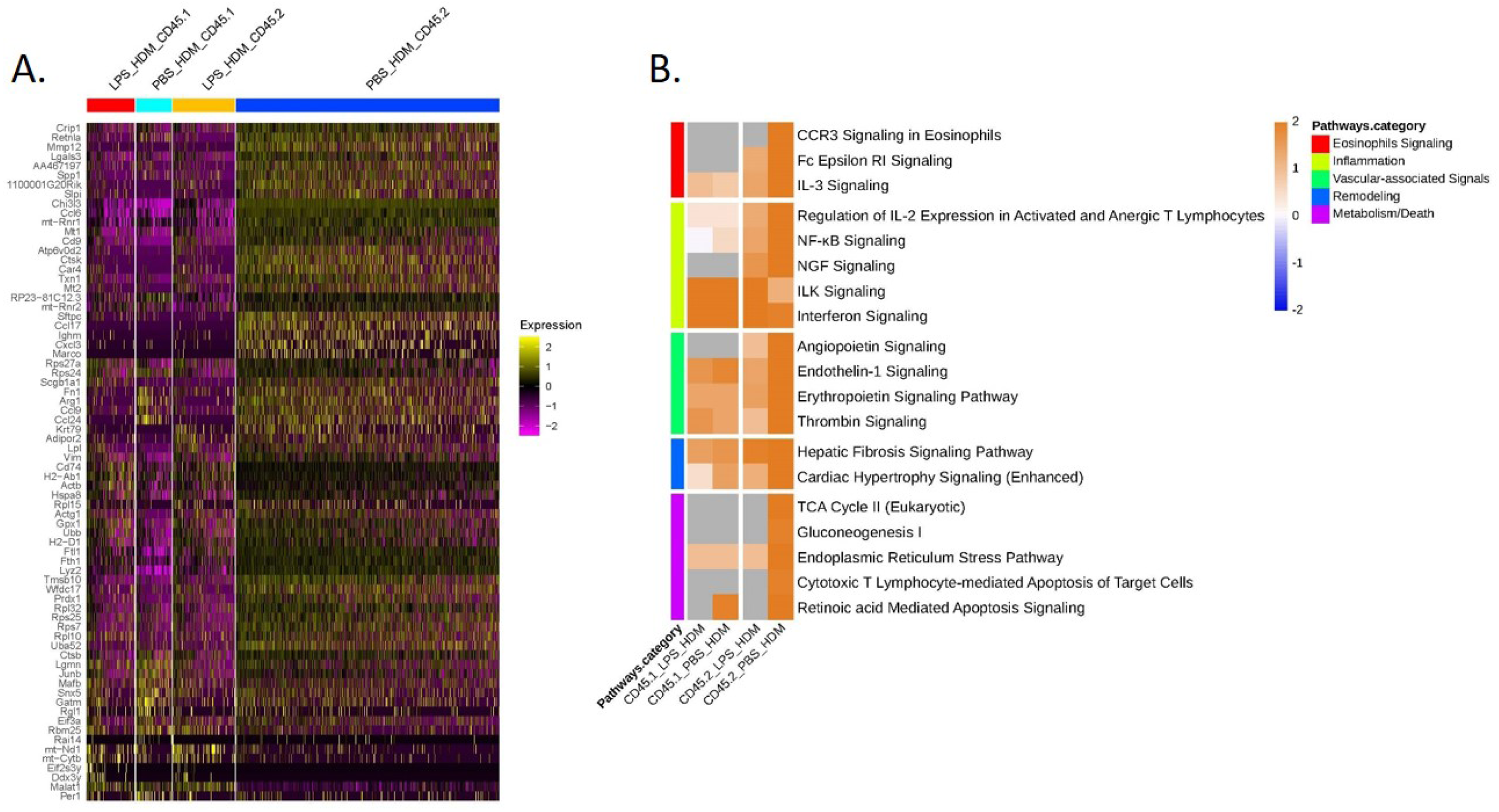
Tissue-resident interstitial macrophages from IM2 cluster are enriched with allergic genes following HDM exposure and abrogated by LPS exposure prior to HDM. A) Top 72 differentially regulated genes from the IM2 cluster from the following exposure conditions and macrophage origin: LPS_HDM in CD45.1 cells (LPS_HDM_CD45.1, red bar), PBS_HDM in CD45.1 cells (PBS_HDM_CD45.1, light blue bar), LPS_HDM in CD45.2 cells (LPS_HDM_CD45.2, orange bar) and PBS_HDM in CD45.2 cells (PBS_HDM_CD45.2, blue bar) expressed as a heatmap. B) Ingenuity pathway analysis (Z score >2 or <2) of IM2 from CD45.1 (recruited) and CD45.2 cells (tissue-resident) based on exposure condition reveals that the IM2 cluster in CD45.2 (tissue-resident) cells is enriched for eosinophilic signaling (red bar), inflammation (yellow bar), vascular-associated signaling (green bar), remodeling (blue bar) and metabolism/cell death (purple bar). In the LPS_HDM exposed CD45.2 cluster, these pathways are reduced compared to the PBS_HDM group.

### IM2 macrophages co-express CD206 and CTSK and are located in the terminal airway parenchyma of PBS-HDM-treated animals

Given the identification of a unique IMØ cluster, which upregulated genes and pathways associated with HDM pulmonary allergic responses, we then attempted to define their location in lung tissue. By assessing top gene expression of individual clusters based on exposure and ontogeny (Figure S7), we identified *Cd206* (a pan-macrophage marker) and *Ctsk* as uniquely co-expressed genes in the IM2 cluster (Figure 7A and B). Immunofluorescence analysis of lung tissue sections revealed a unique population of CD206^+^CTSK^+^ cells located in the parenchyma of the terminal airway in the lungs of PBS_HDM treated mice (Figure 7C and Figure S8). Consistent with our observations in ScRNAseq where, in PBS_HDM treated mice, a unique population of CD45.2 cells was evident and enriched for gene and pathways associated with allergic responses, the CD206^+^CTSK^+^ macrophages were only observed in lung sections of PBS_HDM treated animals. Furthermore, CD206^+^CTSK^+^ double-positive cells were not observed in other conditions (Figure 7C, PBS_PBS, LPS_PBS, or LPS_HDM). This observation supports tissue locational evidence of a unique IMØ population following HDM exposure, which is central to regulating allergic airway responses.

**Figure 7.**
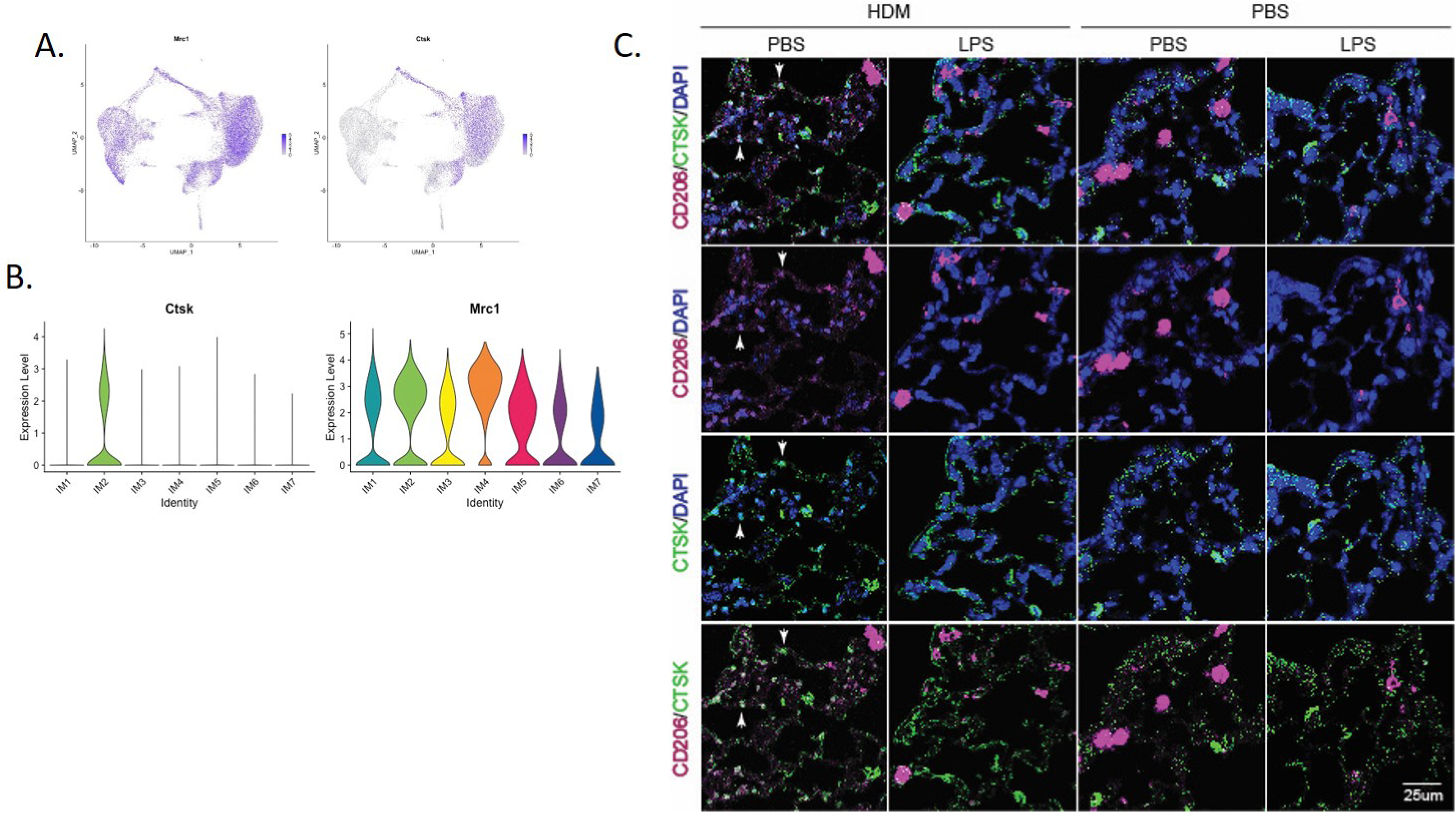
Immunofluorescence staining of IM2 cluster markers defines a subset of interstitial macrophages that localizes to the lung parenchyma adjacent to terminal airways in PBS_HDM exposed mice. A) Evaluation of gene expression patterns across macrophage clusters for pan-macrophage markers revealed broad expression of CD206 (mrc1). Cathepsin K (Ctsk) was widely expressed in alveolar macrophages but in interstitital macrophages was identified as a marker largely restricted to the CD45.2 IM2 cluster in the PBS_HDM exposed mice. B) This was confirmed with violin plots of the IM clusters identifying broad Mrc1 expression but Ctsk expression unique to IM2 cluster C) Immunofluorescence staining on lung tissue sections from PBS_PBS, PBS_HDM, LPS_PBS and LPS_HDM exposed was performed for CD206 (pink, pan-macrophage marker), DAPI (blue, to identify nuclei) and Ctsk (green, CD45.2 IM2 cluster). CD206 staining was noted in macrophages both in the airspace and lung parenchyma in all of the exposure groups. In the PBS_HDM mice, CD206^+^ Ctsk^+^ and DAPI^+^ (white stained cells identified by arrow heads) cells were located in the lung parenchyma adjacent to terminal airways consistent with resident IM2 cluster cells. Co-localization of CD206 and Ctsk in macrophages was not appreciated in the PBS_PBS, LPS_PBS or LPS_HDM groups supporting that these macrophages were unique to the HDM exposed groups.

## Discussion

The present study sought to address if alteration of lung macrophage ontogeny regulates subsequent immune responses in allergic asthma. To address this question, we first demonstrated that the ontogeny of alveolar and interstitial macrophages is distinct, where AMØs appear to arise principally from fetal liver monocytes while the majority of IMØs arise from yolk sac progenitors. Since our focus is on how the changes in the ontological landscape induced by prior lung inflammation affect subsequent immune responses, we developed a model of LPS exposure followed by recovery to homeostasis. Using this model, we demonstrated that LPS exposure induced the replacement of embryonic-derived alveolar and interstitial macrophages with bone marrow-derived alveolar and interstitial macrophages, supporting a change in the ontological landscape upon re-establishment of homeostasis. To assess if this alteration has functional implications for subsequent immune responses, we treated LPS pre-exposed animals (where embryonic-derived macrophages are replaced) with HDM to induce allergic airway disease. We demonstrated that LPS pre-exposed animals were protected from HDM-induced allergic airway disease. To clearly define processes and pathways underlying the protection from experimental allergic asthma and associated with alteration of macrophage ontogenetic landscape, we performed single cell RNA-sequencing from flow-sorted cells based on ontogeny. We observed a specific subset arose from tissue-resident IMØs, located adjacent to terminal airways, and only present in HDM treated without the LPS pre-exposure. This IMØ subset had increased gene expression related to the generation of allergic airway responses. Overall, this data supports that the ontogeny of macrophages, particularly IMØs, is important for allergic airway disease and suggests that approaches to alter macrophage ontogeny may be relevant as a treatment for asthma.

Our focus on ontogeny was due to the observation that lung macrophages exhibit different ontogenies (*i.e.,* fetal-derived or bone marrow-derived). Historically, it was assumed that lung macrophages were either derived from bone marrow origin or largely replaced over the lifespan by bone marrow precursors, thereby limiting the consideration of tissue-resident ontogeny as an effect modifier in immune responses. Recent studies have challenged this notion using detailed lineage tracing and parabiosis system demonstrating, under homeostatic conditions, AMØs are embryonic-derived and exhibit self-renewal. Furthermore, these AMØs appear to arise from fetal liver monocyte precursors (12, 54). IMØ ontogeny has also been defined, though with less overall clarity. Early studies suggest IMØs are yolk sac-derived cells arising around E8.5 or earlier (42, 43). Using a *runx1* reporter to define extra-embryonic yolk sac-derived cells, Tan and colleagues (14) described two populations of macrophages in the interstitium during the early post-natal period: primitive yolk sac-derived and bone marrow-derived. They observed that bone marrow-derived intermediates were gradually recruited to the interstitial space and increased in number over the early post-natal period. In these studies, it was unclear if the bone marrow-derived macrophages eventually overtake yolk sac-derived macrophages as animal age. Subsequently, in adult murine lungs, Chakarov et al. used single cell RNA-seq and S100A4-EYFP lineage labeling to identify two monocyte-derived IMØ populations (defined by differential expression of Lyve1 and MHCII) located either around nerve endings or the vasculature (5). Though they tracked bone marrow-derived IMØ using a S100A4-EYFP reporter, they did not examine the presence of macrophages of embryonic origins. Alternatively, Gibbings et al. identified three distinct IMØ populations by flow cytometry (based on differential expression of Lyve1, MHCII, and CCR2) and demonstrated that these IMØ had differing gene expression (41). This observation was confirmed by Dick et al., using single cell transcriptomic analyses followed by lineage tracing, describing three populations of tissue-resident pulmonary IMØs (16). In contrast to Chakarov et al., they defined a Lyve1+ population arising from both yolk sac and fetal liver monocytes, a MHCII+ population from mixed origins, and a CCR2+ population that is continuously replenished from circulating monocytes. The findings in the present study are largely consistent with those of Dick et al., though we observed that IMØ were mainly yolk-sac derived, without clear derivation from fetal liver monocytes. This discrepancy may have to do with the timing of our lineage induction at E7.5 and E13.5 to distinguish yolk-sac vs. fetal monocyte origins, as opposed to E14.5 and E19.5 in other reports where distinct embryonic origins cannot be distinguished using Cx3cr1^cre/ER^-based lineage tracing system. In addition, we demonstrated that the lineage labeled macrophages were maintained through adulthood, supporting that, under homeostatic conditions, these embryonic-derived IMØs are maintained in the adult. Therefore, our study added to the prior literature by defining distinct ontogenies of lung macrophages preserved into adulthood in the unchallenged state.

Despite defining macrophage ontogeny at steady state in the uninjured mouse, less is understood about how ontogenetic landscape changes with prior lung injury, and how these alterations impact subsequent immune responses. Previous research has focused on differential effects of ontogeny following acute lung injury, particularly differences between tissue-resident and monocyte-derived (i.e., bone marrow-derived) AMØs (3). In models of acute lung injury with bleomycin, asbestosis, influenza, and LPS (9-11, 22-24, 55), bone marrow-derived AMØ are recruited to the lung to direct inflammatory and/or fibrotic responses. In cases of severe lung injury, these bone marrow-derived AMØs can persist after the resolution of lung injury, becoming chimeric with tissue-resident AMØs. Over time and without additional injury, these bone marrow-derived AMØ assume a genetic profile similar to tissue-resident AMØs (10, 11, 55). This process likely reflects an “empty” niche, generated after severe lung injury, where monocyte-derived AMØ can take up residence. Consistent with this, repopulation with yolk-sac, fetal monocyte, or bone-marrow progenitors into Csf2rb-/-mice, which have an empty developmental “niche” for alveolar macrophages, leads to AMØs with nearly identical gene expression patterns independent of the progenitor source (15).

Though the effects of ontogeny in acute lung injury have been defined, less is known about how altered ontogeny from prior injury events that have returned to homeostasis impacts subsequent response. This is particularly relevant as prior exposures or events during lung development or post-natally can change immune cell composition in tissues and thereby have the potential to alter their response to subsequent immune stimuli. Addressing this knowledge gap, we performed LPS exposure to generate lung inflammation and direct turnover of embryonic-derived AMØs and IMØs. We confirmed the return to homeostasis by demonstrating minimal differences in the immune profile through flow cytometry, and cytokine profiles between the PBS_PBS and the LPS_PBS immune cells (Figure 2 and S3). This experimental design was utilized to determine the impact of embryonic-derived lung macrophage replacement without the complicating effect of LPS-induced acute inflammation. When we exposed mice to HDM under conditions where embryonic-derived macrophages had been replaced with monocyte-derived resident AMØ and IMØs, we observed protection from HDM-induced allergic inflammation, mucus hyperplasia, and airway hyperresponsiveness (Figure 2C-F). This data supports that prior exposures can alter subsequent lung responses, and highlights a need for understanding how these prior exposures impact the macrophage compositional profile and function when the lung returns to homeostasis.

Our observations are consistent with recent literature demonstrating that prior murine lung viral infection with murine herpesvirus 4 or murine adapted influenza A attenuates allergic airway inflammation or subsequent pneumovirus infection (56–58). Similar to our observation of AMØ turnover following LPS (Figure 2B), Machiels et al. identified replacement of AMØs following murine herpesvirus 4 infections (56). Furthermore, they reported replacement of AMØs drove the reduced allergic responses. Though our studies also observed replacement of AMØs following LPS exposure, our single cell RNA sequencing studies did not reveal a dominant AMØ cluster in the PBS_HDM exposure groups with upregulated allergic asthma-associated genes (Figure 4 and S5). We also did not observe strong effects of ontogeny on AMØ transcriptomic profile following HDM exposure (CD45.1 vs CD45.2). Despite a lack of an HDM effect, prior exposure to LPS does significantly alter AMØ transcriptomic profile, which could suggest the potential for altered subsequent AMØ responses to immune stimuli that could be evaluated in future studies.

Despite an observed impact on AMØs, Machiels et al. did not explore the impact on IMØs, whose origin is also altered by respiratory viral infections (59) and can impact allergic airway responses (6, 26, 60). Consistent with these IMØ-focused studies, our single cell RNA sequencing studies identified a population of embryonic-derived IMØs (IM2 cluster, Figure 6) that upregulated pathways and processes associated with allergic asthma responses. In addition, using the gene expression data from this cluster, we defined the tissue location of these cells within the lung interstitial spaces adjacent to terminal airways (Figure 7B). This data suggests that embryonic-derived IMØs promote allergic airway responses, and replacing these embryonic-derived IMØs with bone marrow-derived IMØs reduces allergic responses. In addition to the observed replacement of embryonic-derived IMØs with bone marrow-derived IMØs, we also observed an impact of the prior LPS-induced inflammation on the programming of embryonic-derived IMØs, consistent with the concept of trained immunity (61). This conclusion was evidenced in the IM2 cluster from the LPS_HDM_CD45.2 exposure conditions where we observed reduced allergic programming relative to the IM2 cluster from the PBS_HDM_CD45.2 exposure conditions. Overall, our data suggest complex dynamics interplay that drives allergic airway disease, involving interstitial macrophage ontogeny and trained immunity by prior lung injury.

To our knowledge, this study is the first description of how prior exposures shapes the subsequent ontogenetic landscape of pulmonary macrophages and their functional profile. The observation has important implications for understanding human lung immunity. Humans do not exist in a sterile environment but experience environmental challenges over their lifespan that can impact allergic responses. Consistent with this observation, prior environmental LPS exposure has been associated with reduced allergic sensitization and asthma (62, 63). Our data support that IMØs may exert a central role in these responses. Therefore, understanding the dynamics of prior exposure to macrophage functions in homeostasis and response to subsequent challenges is critical. Though not explicitly addressed in this initial study, future studies will be required to assess the differential impact of replacement versus memory of prior exposures on allergic airway responses.

Some limitations should be considered in the present study. Though the focus of our study was on macrophages, we acknowledge that T cells are an essential effect modifier of allergic airway responses (64), they exhibit cross-talk with macrophages in this context (65) and their function can be altered by LPS exposure. T cell-specific effects on allergic responses will need to be considered in future studies and analyses. In addition, we demonstrated that the IM2 cluster upregulates gene related to allergic airway responses and are located in the tissue adjacent to terminal airway structures. However, as the methods for the targeted examination of this specific cluster are not presently available, we could not demonstrate clear causality for this IMØ subset in promoting allergic responses. Despite efforts to ablate IMØ by chemical or genetic methods, we were not able to target IMØ with specificity or without significant associated inflammation (data not shown). Future efforts will need to identify targeted strategies that impact IMØ subsets for causal studies.

In summary, we defined distinct ontogenies of AMØ and IMØ. Their embryonic ontogeny is maintained in adult mice but can be altered by prior LPS exposure. LPS exposure attenuates subsequent HDM-mediated allergic airway responses. This effect is mediated by IMØs via alteration in ontogenetic composition and change in genetic programming. Overall, this study supports that lung macrophage ontogeny is important for allergic airway responses and highlights a role for IMØ ontogeny in these responses.

## Abbreviations

AHR: Airway hyper-responsiveness
AMØ: Alveolar macrophage
IMØ: Interstitial macrophage
T2: T helper type 2
HDM: House dust mite
o.p.: Oropharyngeal aspiration
MCh: Acetyl-β-methacholine
R_rs_: Respiratory system resistance
E_rs_: Respiratory system elastance
Rn: Newtonian resistance
G: Tissue damping
H: Tissue elastance
BALF: Bronchoalveolar lavage fluid

## Acknowledgments

The authors thank the Duke University Rodent Inhalation Core Facility for assisting with mouse exposures and airway physiology measurements.

