## Supplementary material for "Allergic Asthma Responses Are Dependent on Macrophage Ontogeny": Online supplement

Table S1 - Flow Cytometry Antibodies in base panel and additional markers to confirm cell type

| Antibody | Clone | Dilutions | Fluorochrome | Company | Catalog # |
|---|---|---|---|---|---|
| CD11b | M1/70 | 1:50 | APC-Cy7 | BD Biosciences | 557657 |
| CD11c | HL3 | 1:100 | BV785 | BD Biosciences | 563735 |
| CD24 | M1/69 | 1:800 | BV711 | BD Biosciences | 563450 |
| CD45 | 30-F11 | 1:500 | BV605 | Biolegend | 103139 |
| CD49b | DX5 | 1:100 | PE | eBioscience | 12-5971-83 |
| CD64 | X54-5/7.1 | 1:200 | BV421 | Biolegend | 139309 |
| IA/IE | M5/114.15.2 | 1:1500 | BV650 | BD Biosciences | 563415 |
| Ly6C | HK1.4 | 1:200 | PerCP-Cy5.5 | eBioscience | 45-5932 |
| Ly6G | 1A8 | 1:200 | AF700 | BD Biosciences | 561236 |
| Zombie Yellow | NA | 1:500 | ex:405nm/em:572nm | Biolegend | 423104 |
| SiglecF | E50-2440 | 1:500 | PE-CF594 | BD Biosciences | 562757 |
| CD3 | 145-2C11 | 1:100 | AF488 | Biolegend | 100321 |
| CD31 | MEC 13.3 | 1:400 | PerCP-Cy5.5 | Biolegend | 562861 |
| CD117 | 2B8 | 1:100 | BV510 | Biolegend | 105839 |
| B220 | RA3-6B12 | 1:100 | AF647 | Biolegend | 103226 |

| | | | | | |
|---|---|---|---|---|---|
| F4/80 | BM8 | 1:400 | PE-Cy7 | eBioscience | 25-4801 |
| FceR1 | MAR-1 | 1:100 | PE-eF610 | eBioscience | 61-5898-82 |

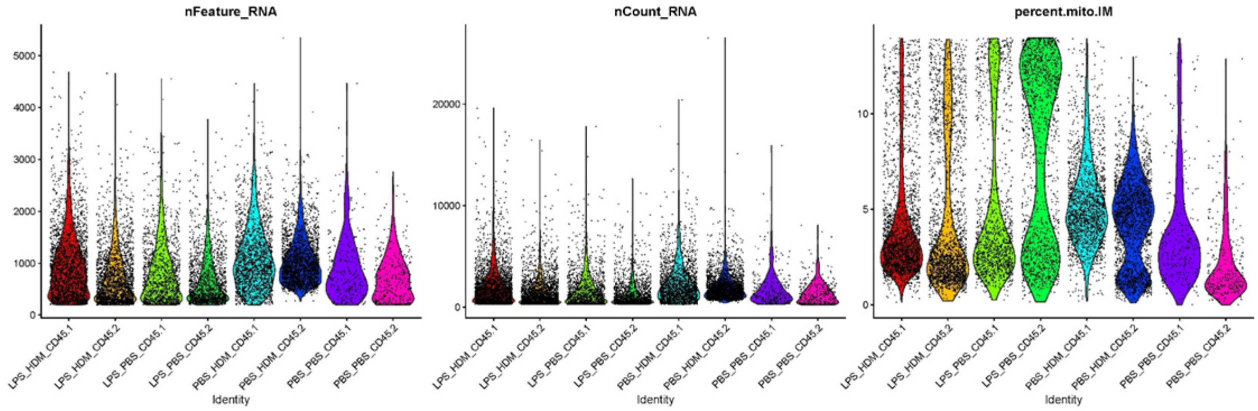

**Figure S1**: *Increased mitochondrial gene cutoff from initial IM clustering to remove cell cluster artifact of damaged cells.* Interstitial macrophage clustering was assessed for expression of mitochondrial genes. A) During initial clustering evidence of high mitochondrial gene expression was appreciated in one of the clusters of interest. B) This was also considered by violin pilots based on the different cell source (recruited versus resident derived) and exposure to define an improved cutoff for mitochondrial genes.

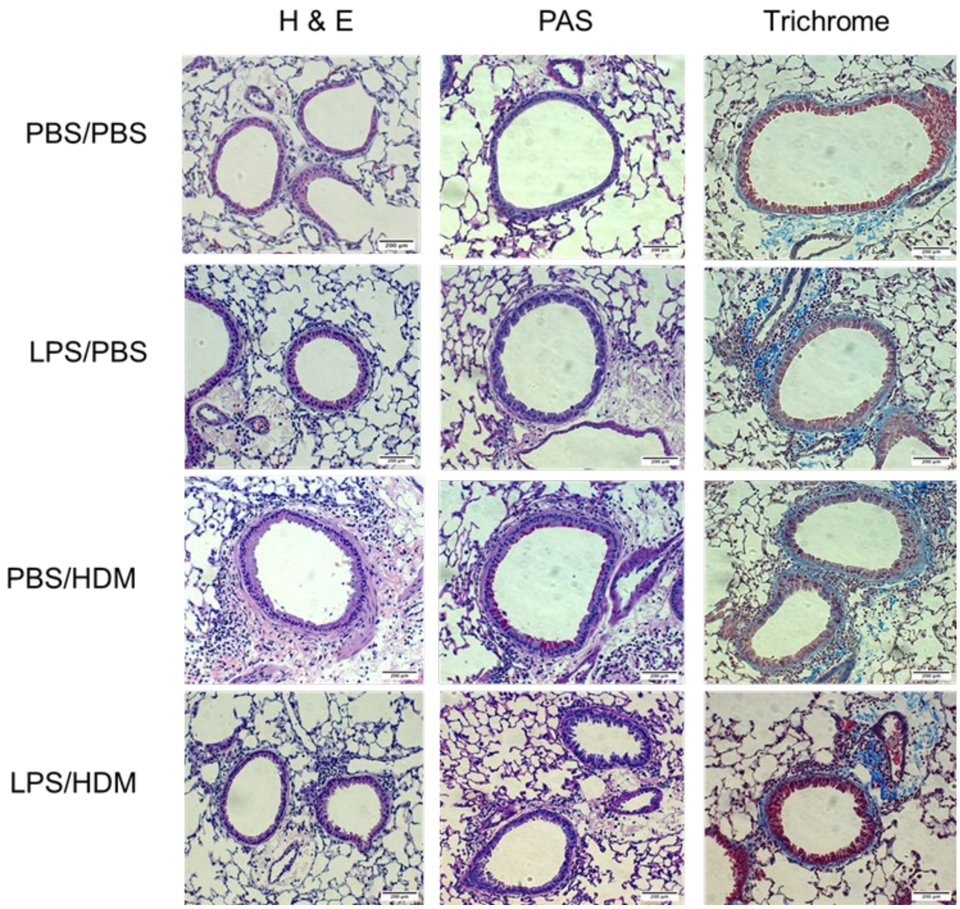

**Figure S2**: *Representative images for H&E, Trichrome and PAS staining from the PBS/PBS, LPS/PBS, PBS/HDM and LPS/HDM groups.* Left lung at the time of necropsy was isolated and inflated with formalin to fix the lung tissue. The tissue was then processed and sectioned. Staining was performed for hematoxylin and eosin (H & E) to assess inflammation, Periodic acid–Schiff (PAS) to assess for mucins, and trichrome to assess for evidence of fibrosis. Images are representative of sections from each exposure condition with a focus on similar regions (larger airways) of the lung tissue. Scale bar represents 200 μm.

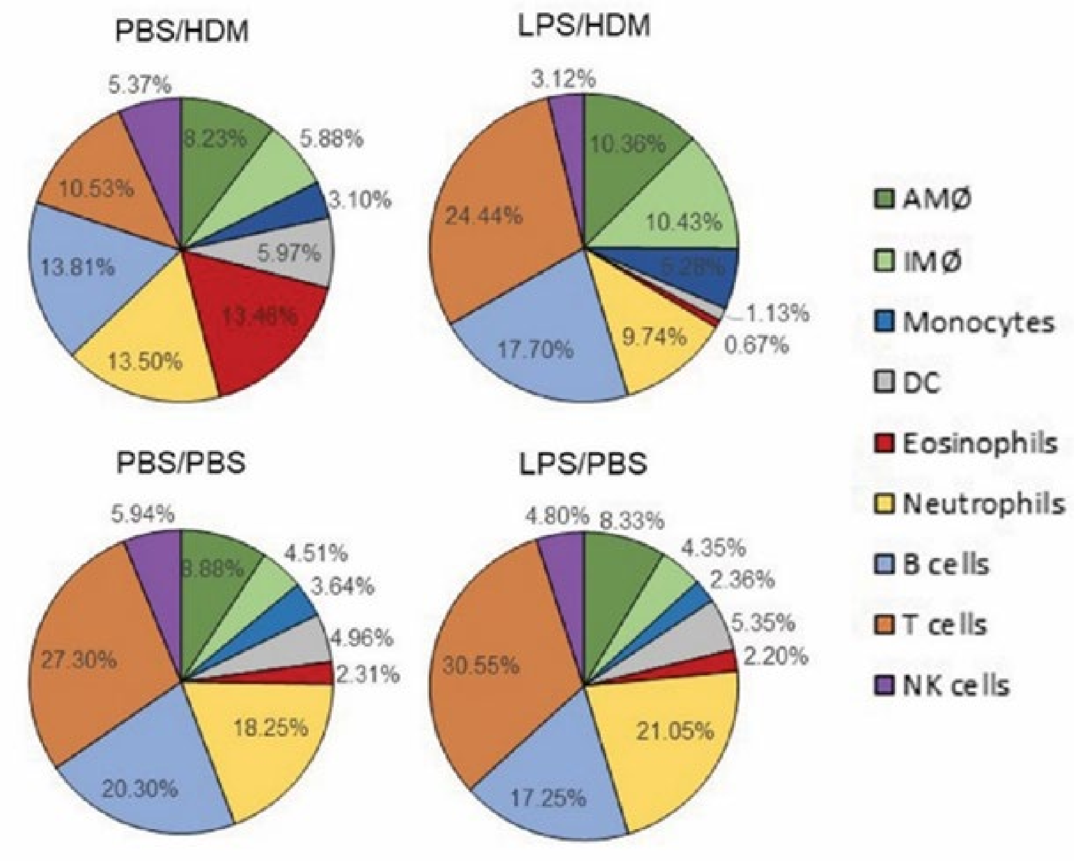

**Figure S3**: *Lung tissue immune cell composition by flow cytometry is dependent on exposure condition*. Cx3cr1cre-ER;TdTomato mice underwent lineage reporter induction with tamoxifen at E13.5 and then allowed to age till 6 weeks. At 6w, i.n. LPS or PBS exposure occurred and then the mice were allowed to recover for 8 weeks. Following recovery mice were exposed to PBS or HDM once per week for 3 weeks and then lung tissue was harvested and processed for flow cytometry 48h after the last exposure. Using an established flow cytometry panel that identifies the principal immune cell components, we define alveolar macrophages (AMØ), interstitial macrophages (IMØ), monocytes, dendritic cells (DC), eosinophils, neutrophils, B cells, T cells and natural killer (NK) cells. Values are defined as a percentage of live, CD45+ cells for the following exposure conditions: 1) PBS followed by HDM (PBS/HDM), 2)

LPS followed by HDM (LPS/HDM), 3) PBS followed by PBS (PBS/PBS), and 4) LPS followed by PBS (LPS/PBS). N=5 samples per exposure condition.

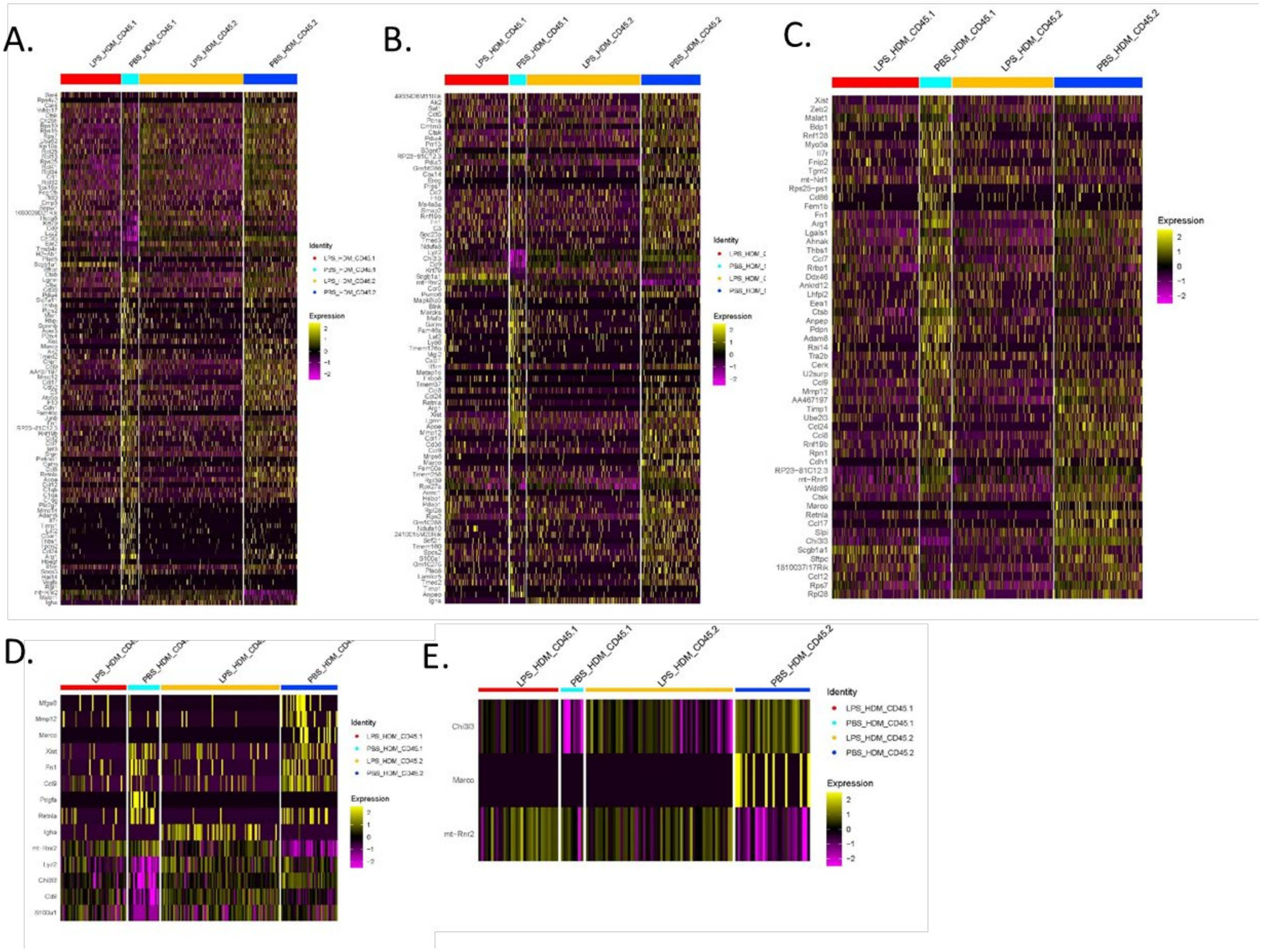

**Figure S4**: *Heat map of gene expression for AMØ clusters.* Top differentially expressed genes were defined for each AMØ cluster and then were expressed as a heat map segregated base on the following conditions LPS_HDM in CD45.1 cells (LPS_HDM_CD45.1, red bar), PBS_HDM in CD45.1 cells (PBS_HDM_CD45.1, light blue bar), LPS_HDM in CD45.2 cells (LPS_HDM_CD45.2, orange bar) and PBS_HDM in CD45.2 cells (PBS_HDM_CD45.2, blue bar). This allowed comparison of gene expression patterns between exposure condition and based on cell origin in AM1 (A), AM2 (B), AM3 (C), AM4 (D), and AM5 (E) clusters.

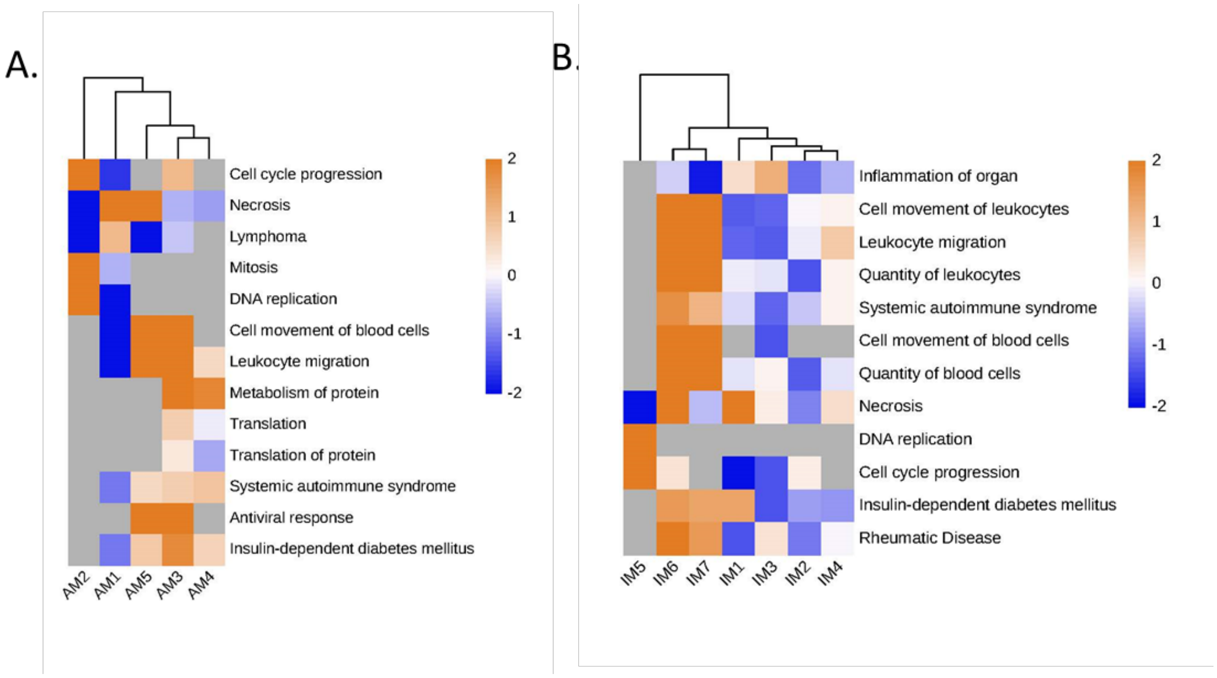

**Figure S5**: *Pathway analysis per overall cluster in AMs and IMs, independent of exposure condition.* Ingenuity pathway analysis was performed on each alveolar macrophage (AM, A) and interstitial macrophage (IM, B) cluster independent of ontogeny or exposure condition to define overall patterns of the individual clusters. Hierarchical clustering was performed to define potential associations.

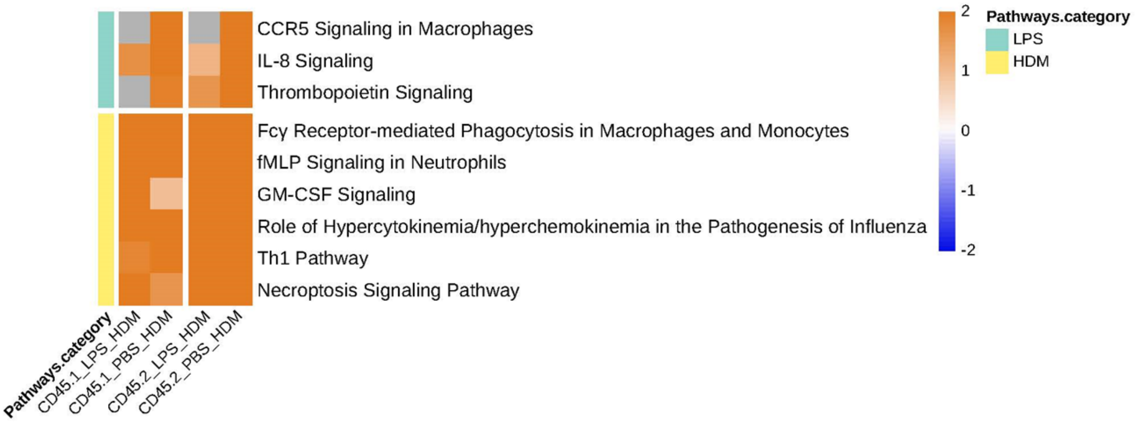

**Figure S6**: *Pathway analysis in CD45.1 and .2 IM cluster 2 cells based on exposure condition compared to PBS-PBS groups with a focus on common pathways in LPS and HDM responses across the exposure conditions.*

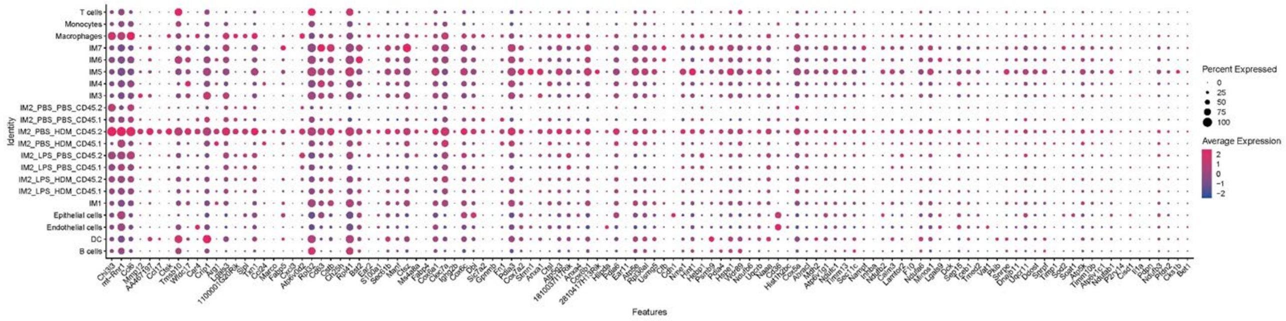

**Figure S7**: *Gene analysis of all immune cells, interstitial macrophages and the IM2 cluster based on exposure conditions to define specific markers for the IM2 cluster to use to identify tissue location.* Differential gene expression was performed to define potential markers that uniquely identified the IM2 cluster to consider for tissue immunofluorescence staining.

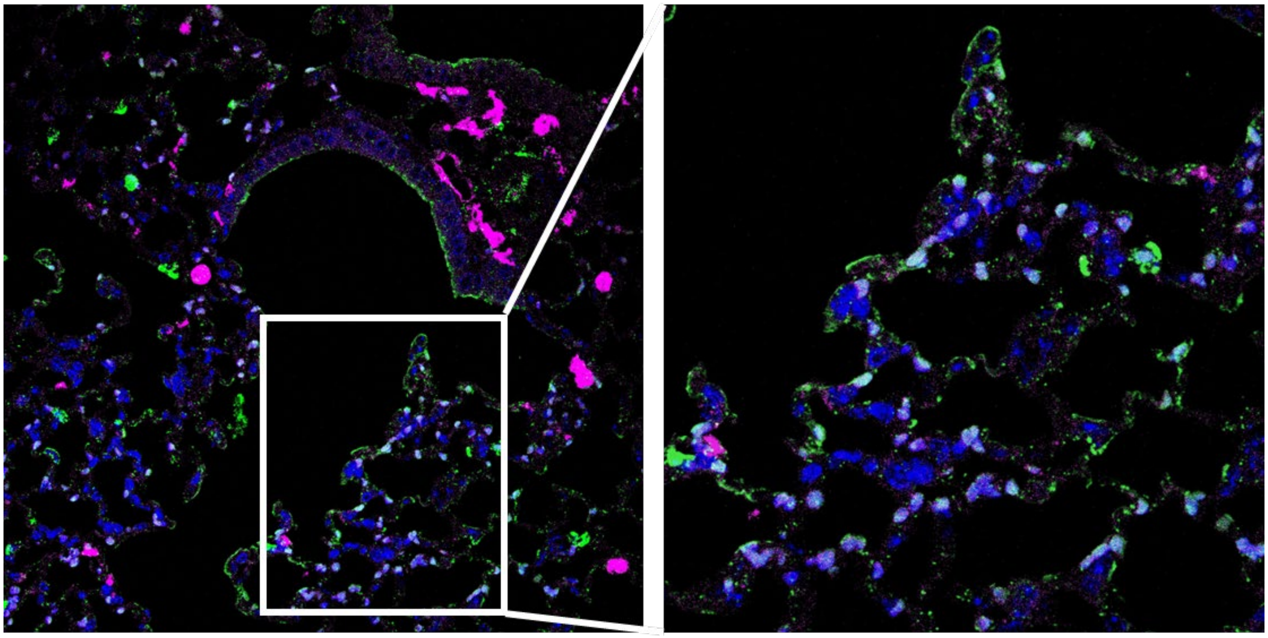

**Figure S8**: *Enlarged images of immunofluorescence staining for CD206 and Ctsk to define the IM2 cluster in lung tissue.* CD206 (pink, pan-macrophage marker), DAPI (blue, to identify nuclei) and Ctsk (green, CD45.2 IM2 cluster).
